# The link between steady-state EEG and rs-fMRI metrics in healthy young adults: the effect of macrovascular correction

**DOI:** 10.1101/2025.06.06.658306

**Authors:** Xiaole Z. Zhong, Hannah Van Lankveld, Alicia Mathew, J. Jean Chen

## Abstract

To improve the clinical utility of resting-state fMRI (rs-fMRI), enhancing its interpretability is paramount. Establishing links with electrophysiological activities remains the benchmark for understanding the neuronal basis of rs-fMRI signals. Existing research, while informative, suffers from inconsistencies and a limited scope of rs-fMRI metrics (e.g., seed-based functional connectivity). Phenotypic variables like sex and age are suspected to obscure reliable fMRI-EEG associations. A major contributing factor to these inconsistencies may be the neglect of macrovascular correction in rs-fMRI metrics. Given that macrovascular contributions can inflate rs-fMRI connectivity and power, they may lead to misleading fMRI-EEG associations that do not reflect genuine neuronal underpinnings. In this study, we addressed this by applying macrovascular correction and performing a systematic, inter-participant analysis of multiple rs-fMRI and EEG metrics. Our key findings demonstrate that: 1) Macrovascular correction enhances the relationship between EEG and rs-fMRI metrics and improves model fit in many instances; 2) sex significantly modulates EEG-fMRI associations; 3) EEG complexity is significantly associated with resting-state functional activity (RSFA). This research provides crucial insights into the interplay between rs-fMRI and EEG, ultimately improving the interpretability of rs-fMRI measurements and building upon our prior work linking fMRI and metabolism.

## 1 Introduction

Resting-state functional magnetic resonance imaging (rs-fMRI) using blood-oxygenation level-dependent contrast (BOLD) has been widely used to understand brain function (Beckmann et al., 2005). The measurements of rs-fMRI can be interpreted in accordance with their connectivity patterns (Biswal et al., 1995), temporal variability (Tsvetanov et al., 2015; Zhong & Chen, 2022), and temporal complexity (Z. Wang, 2021; Zhong et al., 2025). Nevertheless, promoting the clinical application of rs-fMRI is challenging due to the complexity of the BOLD signal. Our previous study established inter-participant associations of rs-fMRI measurements with cerebral oxidative metabolism (CMRO_2_) and its correlates, which provide a healthy-adult reference for metabolism underpins of rs-fMRI. Despite being closer to neuronal activity than BOLD-based rs-fMRI, oxidative metabolism does not fully reflect localized neuronal activity, as discussed in our previous study (Zhong et al., 2025). Thus, it is still considered necessary to link rs-fMRI measurements to electrophysiological activity. Nevertheless, associations established by most previous studies have been mainly based on inter-regional differences, which are strongly affected by conditions (Wirsich et al., 2021) and phenotypes (Nentwich et al., 2020), leading to a lack of generalizability and consistency across studies. Hence, establishing the inter-participant electrophysiology-fMRI associations that mirror our CMRO_2_-fMRI associations could provide a more generalizable and consistent for understanding neuronal underpinnings of rs-fMRI measurements.

The most direct approach to understanding the electrophysiological underpinnings of rs-fMRI (if not stated otherwise, ‘rs-fMRI’ in this paper would refer to GRE-BOLD-based rs-fMRI) is to correlate fMRI metrics with electroencephalography (EEG) metrics. In principle, neuronal connectivity measured by EEG and rs-fMRI functional connectivity (FC) should be at least partially associated with one another, given that both modalities have the same neuronal basis for their recording (Tu et al., 2024). This is indeed observed in some human experiments, but findings have not been consistent. Wirsich et al. reported that the strongest associations (inter-regional) were found between the rs-fMRI FC and the EEG FC in the beta and alpha bands (Wirsich et al., 2021). Similar results were also obtained by an experiment with monkeys, indicating that interareal connectivity within the visual cortex is dominated by low frequency coherence (< 20 Hz) in the eyes-closed condition (L. Wang et al., 2012). On the other hand, based on a comparison of spatial extents, Deligianni et al. found EEG FC based on the delta and theta bands to exhibit the strongest association with rs-fMRI FC (Deligianni et al., 2014). Moreover, EEG entropy was found to be negatively associated with EEG FC in the mouse model (M. Liu et al., 2019), therefore, it would be reasonable to expect that there would be a negative association between EEG signal complexity and rs-fMRI FC. However, there are also other studies that report weak or even no correlation between EEG FC and rs-fMRI FC (Nentwich et al., 2020; Wirsich, Amico, et al., 2020).

There have also been modulatory effects observed between EEG power and seed-based rs-fMRI FC, where higher alpha power is associated with a lower visual cortex rs-fMRI FC strength (Scheeringa et al., 2012) but a larger spatial extent for the primary visual network (Phadikar et al., 2025). Similar modulatory effects may extend to other EEG bands and networks (Z. Liu et al., 2014; Neuner et al., 2014). The EEG-modulatory effects are also evident in terms of dynamic seed-based rs-fMRI FC (Chang et al., 2013; Tagliazucchi et al., 2012), but not dominated by power or connectivity from any specific EEG band (Wirsich, Giraud, et al., 2020). Thus, the association between rs-fMRI metrics and broadband EEG patterns may also be informative of neuronal underpinnings of rs-fMRI FC. Higher spontaneous fluctuations in neuronal activity can be associated with a higher oxidative metabolism (Hudetz et al., 1998), which is in turn associated with higher rs-fMRI FC (inter-participant), especially functional-connectivity density (FCD), as shown by our previous study (Zhong et al., 2025). Taken together, it is reasonable to hypothesize a systematic link between EEG power and rs-fMRI FC due to their respective links to metabolism. Moreover, most previous studies have used seed-based BOLD connectivity (Nentwich et al., 2020; L. Wang et al., 2012). Thus, to better understand the links between EEG and fMRI connectivity, a broader range of metrics (e.g. FCD) may be required.

Given the dependence of rs-fMRI FC on BOLD signal amplitude (Lee et al., 2023), it is reasonable to expect an association between rs-fMRI signal power and EEG signal power (Hyder & Rothman, 2010). If we assume that resting-state fluctuation amplitude (RSFA) (as rs-fMRI power) is driven by the same neurovascular principles as task-evoked BOLD (Logothetis et al., 2001), then we can hypothesize that, based on past task-based studies (Guy et al., 1999; Lachaux et al., 2007), that the alpha and gamma EEG power are most strongly associated with BOLD RSFA. However, most such studies were based solely on the visual or default-mode regions, which may not generalize to the rest of the brain. Liu et al. found that all EEG bands contribute to spontaneous fluctuations in BOLD (Z. Liu et al., 2014). Moreover, the assumption of neurovascular coupling may not hold in the resting state. Indeed, there are limited associations found between the inter-participant difference in RSFA and EEG power (Kumral et al., 2020; Tsvetanov et al., 2015) or EEG frequency (Zhong & Chen, 2022). Thus, it has been suggested that RSFA is more closely associated with vascular activity than neuronal activity, especially in the brain-aging context (Tsvetanov et al., 2020). Nevertheless, our previous study suggests that EEG complexity index (CI, an entropy-based metric) may partially mediate the age effect on RSFA, which raises the possibility of EEG complexity being associated with the RSFA (Zhong & Chen, 2022).

The concept of entropy has also become adopted in rs-fMRI. Compared to linear metrics such as FC and RSFA, rs-fMRI entropy measurement can provide additional insight into spontaneous brain activity (Nezafati et al., 2020; Saxe et al., 2018; Song et al., 2019; Xin et al., 2024). However, as a newly developed metric for rs-fMRI, the biophysical and physiological mechanisms behind the rs-fMRI entropy remain unclear. Investigating the associations between rs-fMRI entropy and EEG metrics can help us clarify these mechanisms. In our previous research, we demonstrated that EEG complexity provided the only link between rs-fMRI and EEG variations in aging (Zhong & Chen, 2022), with both rs-fMRI entropy and EEG complexity becoming reduced in older adults (Del Mauro et al., 2024; Zhong & Chen, 2022). Hence, rs-fMRI entropy should be positively associated with the EEG complexity outside the aging context, if indeed they reflect the same neuronal mechanisms. There is, however, no established link between rs-fMRI entropy and other EEG metrics, and whether there are meaningful neuronal interpretations for rs-fMRI entropy remains an important unanswered question. The results of our previous study (Zhong et al., 2025) revealed a significant inter-participant relationship between rs-fMRI entropy and oxidative metabolism, suggesting possible associations between rs-fMRI entropy and EEG metrics due to their respective links to metabolism.

One source of discrepancies between EEG and rs-fMRI measurements may be macrovascular bias in rs-fMRI, which obscures the neuronal contributions to rs-fMRI. Our recent study (Zhong, Tong, et al., 2024) suggests that macrovascular bias (especially from veins) may extend to perivascular tissues extending as far as 1.2 cm from the boundary of the blood vessel. As a consequence, the local-neuronal specificity of rs-fMRI could be lost, and links between these measurements and EEG measurements may become inconsistent. Nevertheless, our recent findings also indicate that the macrovascular bias can be biophysically modelled (Zhong, Polimeni, et al., 2024). Using a biophysical-informed modeling-based correction procedure, we demonstrate improvements in the extent to which rs-fMRI metrics are explained by cerebral metabolic rate of oxygen (Zhong et al., 2025). This newly introduced method could also facilitate the investigation of associations between EEG and rs-fMRI in the present study.

Many of the previous studies (as listed above) examine dynamic associations between rs-fMRI FC and EEG FC over a relatively short time. These associations, however, require simultaneous EEG-fMRI recordings, which present their own challenges in acquisition and analysis, and are not typically acquired in large numbers. Alternatively, establishing the associations between static (averaged over time) EEG and rs-fMRI metrics (Nentwich et al., 2020; Wirsich et al., 2021; Zhong & Chen, 2022) can help elucidate the EEG-fMRI associations without simultaneous recordings. Moreover, while most past studies have focused on inter-regional and dynamic associations between EEG and fMRI, these generalizability of these associations may be hampered by differences not only across conditions (e.g. eyes-open vs. eyes-close) (Wirsich et al., 2021), but also across participants (e.g. sex and age) (Nentwich et al., 2020). More broadly, such associations can also differ between healthy and patient cohorts. Thus, examining the EEG-fMRI associations driven by differences across participants offers a way to capture consistencies in the EEG-fMRI associations across individuals and conditions that could be helpful in establishing a more generalizable physiological reference point. By establishing such a reference, one may be able to detect abnormalities in aging and disease. In this work, we build on previous studies, including our own, but distinct from previous work, we base our conclusions on rs-fMRI metrics corrected for macrovascular bias (Zhong et al., 2025). We hypothesize that (1) macrovascular correction can enhance the extent to which EEG metrics can explain fMRI metrics; (2) associations between metrics of EEG and rs-fMRI will differ by sex, based on previous work highlighting the importance of phenotype <u>(Nentwich et al. 2020; Zhong and Chen 2022; Zhong et</u> <u>al. 2025)</u>; (3) similar to our findings in aging (Zhong and Chen, 2022), fMRI RSFA will be most strongly associated not with EEG power but with EEG complexity. We believe a systematic analysis of the activity of the EEG underpinning common rs-fMRI metrics will improve the interpretability of the rs-fMRI measurements.

## 2 Method

This study used the same methodology for *participant recruitment*, *MRI acquisition*, *macrovascular correction*, and *rs-fMRI processing* as in our previous study (Zhong et al., 2025).

### 2.1 Participant

A total of 20 healthy participants are involved in this study (10 males and 10 females between the ages of 20 and 32 years). No participant reported having a history of cardiovascular disease, psychiatric illness, neurological disorder, malignant disease, or medication use that may have affected the study. Recruiting participants was conducted through the Baycrest Participants Database, which includes residents of Baycrest and surrounding areas. A research ethics board (REB) of Baycrest approved the study, and the experiments were carried out with the understanding and consent of each participant.

### 2.2 MRI acquisition

The images were acquired using a Siemens Prisma 3 Tesla System (Siemens, Erlangen, Germany), which employed 20-channel phased-array head coil reception and body coil transmission. The participants were imaged during naturalistic-stimulus viewing to reduce random mind-wandering which increases reproducibility (Gal et al., 2022).

The following data were acquired for each participant: (i) T1-weighted structural image (sagittal, 234 slices, 0.7 mm isotropic resolution, TE = 2.9 ms, TR = 2240 ms, TI = 1130 ms, flip angle = 10°); (ii) two time-of-flight (TOF) scans, with coronal and sagittal flow encoding, respectively (coronal: 0.8 x 0.8 x 2.125 mm thickness, 100 slices, TR = 16.6 ms, TE = 5.1 ms, flip angle (*α*) = 60°; sagittal: 0.8 x 0.8 x 2.125 mm thickness, 80 slices, TR = 16.6 ms, TE = 5.1 ms; flip angle (*α*) = 60°); (iii) one dual-echo (DE) pseudo-continuous arterial spin labelling (pCASL) (courtesy of Danny J. J. Wang, University of Southern California) for recording cerebral blood flow (CBF) and BOLD dynamics (TR = 4.5 s, TE1 = 9.8 ms, TE1 = 30 ms, post-labelling delay = 1.5 s, labelling duration = 1.5 s, flip angle (*α*) = 90°, 3.5 mm isotropic resolution, 35 slices, slices gap = 25%, scanning time = 4 minutes).

### 2.3 EEG acquisition

In a separate session from the MRI, a 256-channel Geodesic Sensor Net with a Net Amp 400 amplifier (Magstim Inc., Roseville, MN, USA) was used to record 4 minutes of EEG. The participants were instructed to view a naturalistic stimulus video to reduce random mind-wandering. The EEG electrodes were attached following the international standard 10-20 system, and the impedance of each electrode was controlled per the vendor recommendations (lower than 50 kOhm recommended for the saline-based system). The EEG signal was digitized at a sampling frequency of 1,000 Hz, and the amplitude resolution was set to 0.024 μV.

### 2.4 Macrovascular correction

A detailed overview of the BOLD macrovascular correction framework can be found in our previous study (Zhong et al., 2025) and a more detailed description of the related biophysical modelling can be found in another our previous study (Zhong, Polimeni, et al., 2024). Since the scan parameters were the same between this study and the previous study (Zhong et al., 2025) that applied macrovascular correction, none of the parameters in the correction pipeline had been modified.

A summary of the macrovascular correction pipeline is provided below: 1) generate a vascular mask for each participant and estimate the blood-volume fraction (fBV); 2) extract the input signal from the voxel with the max fBV; 3) simulate macrovascular BOLD signals using our macro-VAN model; 4) assess the lag between the simulated signal and the in-vivo signal for each voxel (lag that leads to maximum absolute cross-correlation coefficient); 5) regress the simulated signal from the in-vivo signal for each voxel as nuisance regressor (linear regression model).

### 2.5 fMRI preprocessing and analysis

Two pipelines were used to process the rs-fMRI data: one for investigating seed-based functional connectivity (FC) and the other for seed-independent measures. Both pipelines include preprocessing steps that include rejecting the first five volumes of BOLD data to allow the brain to enter the steady state. Furthermore, all metrics below were averaged within each functional network listed in the *Seed-based rs-fMRI analysis* subsection to calculate metrics at the network level.

#### 2.5.1 Seed-based rs-fMRI analysis

rs-fMRI FC analysis was conducted using the CONN toolbox (Whitfield-Gabrieli & Nieto-Castanon, 2012) with the default_MNI preprocessed pipeline (bandpass filter: 0.01-0.1 Hz; regressed confounds: white matter, CSF and estimated participant-motion parameters). The networks of interest include the visual network (VN), sensorimotor network (SMN), the default mode network (DMN), the language network (LN), the salience network (SN), the dorsal attention network (DAN), and the frontoparietal network (FPN). For each network, p-value maps for different sources were combined by selecting a minimum p-value and then thresholding by p < 0.0001. Network-wise seed-based FC was calculated by averaging the magnitude of correlation coefficients within each network.

#### 2.5.2 Seed-independent rs-fMRI analysis

rs-fMRI preprocessing pipeline was implemented with customized procedures based on tools from FSL (Jenkinson et al., 2012), AFNI (Cox, 1996) and FreeSurfer (Fischl, 2012). The following steps were included in the preprocessing steps: (a) 3D motion correction (FSL MCFLIRT), (b) slice-timing correction (FSL slicetimer), (c) brain extraction (FSL bet2 and FreeSurfer mri_watershed), (d) rigid body coregistration of functional data to the individual T1 image (FSL FLIRT), (e) regression of the mean signals from white-matter (WM) and cerebrospinal fluid (CSF) regions (fsl_glm), (f) bandpass filtering to obtain frequency band 0.01-0.1 Hz (AFNI 3dBandpass), and (g) spatial smoothing with a 6 mm full-width half-maximum (FWHM) Gaussian kernel (FSL fslmaths).

Once the BOLD signal had been preprocessed, we computed the following metrics:

1. Resting-state fluctuation amplitude (RSFA), is calculated using the standard deviation of the BOLD signal normalized by the mean of the BOLD signal.
2. BOLD signal entropy, which quantifies the unpredictability or complexity of BOLD signal fluctuations across time, is calculated using sample entropy with m = 3 and r = 0.6 related to the standard deviation as proposed by the previous study (Nezafati et al., 2020).
3. lFCD, the number of connections within the immediate neighbourhood, calculated using AFNI (3dLFCD) (Cox, 1996) with a threshold of 0.6, and the neighbourhood data was defined to include face-touching 6 voxels, as suggested by the previous study (Tomasi & Volkow, 2010).
4. gFCD, the number of connections with all other regions in the brain, is calculated using the same threshold value (0.6) but within the entire gray matter (Tomasi & Volkow, 2011).

For lFCD and gFCD, log-transformed data were used since density-related measurements follow an exponential distribution (Shokri-Kojori et al., 2019).

### 2.6 EEG preprocessing and analysis

#### 2.6.1 Preprocessing

EEG preprocessing was carried out using EEGLAB (version 2019.1) (Delorme & Makeig, 2004) with functions implemented in Matlab (The MathWorks Inc., Natick, Massachusetts, USA). The following steps were included in the preprocessing pipeline: 1) removal of face electrodes; 2) downsampling to 125 Hz; 3) high-pass filtering with a cut-off frequency of 1 Hz; 4) ICA denoising (removal of components related to physiological sources, including eye blinks, eye movements, residual ballistocardiograph artifacts and muscle activity); 5) channel rejection (criteria: more than 80% noisy data, line noise exceeding 4 standard deviations, flat-line segments of more than 5 seconds); 6) linear detrending.

The source reconstruction was performed using the Cartool toolbox (version 4.11) (Brunet et al., 2011). The forward model was developed for each participant using the T1-weighted MRI scans and the Locally Spherical Model with Anatomical Constraints (LSMAC) model (6-shell). The thickness and conductivity of the skull were adjusted according to each participant’s age. The inverse model was estimated using exact low-resolution brain electromagnetic tomography (eLORETA) (Imperatori et al., 2020) with 6000 solution points. The dominant orientation of the source signal was determined using singular-value decomposition (SVD) of each voxel. In addition, the sources were further mapped back to the fMRI space based on coordinates with the nearest neighbour principle so that the following metrics could be averaged with the same network ROIs as the fMRI metrics.

#### 2.6.2 EEG power

The Fourier transform was applied to the signal from each source. Band-specific fractional power was determined by dividing the power of each band by the total power across all bands, with bands defined as follows: delta (1-4 Hz), theta (4-8 Hz), alpha (8-12 Hz), beta (12-30 Hz) and low gamma (30-50 Hz). This is the same definition of bands as used in our previous study (Zhong & Chen, 2022). Unlike raw band-limited power, normalized power accounts for fluctuations in total power while quantifying the contribution of each band. Additionally, the total power across all bands was normalized by the spatial standard deviation of the total power for each participant to account for different SNRs across participants.

#### 2.6.3 EEG coherence

The imaginary part of EEG-signal coherence has been widely used to measure synchronization between two EEG signals, since it is less sensitive to volume conduction effects (Nolte et al., 2004). Our study utilized this approach to quantify EEG-based brain connectivity (Nentwich et al., 2020; Nolte et al., 2004) with a frequency resolution of 0.5 Hz and an overlap of 0.5 using scripts adapted from the Brainstorm toolbox (Tadel et al., 2011). Band-limited coherence for each band was calculated by averaging the coherence spectrum over the frequency range of each band as defined earlier. Moreover, total coherence was calculated by averaging the coherence spectrum across all bands. Here, band-limited coherence reflects functionally specific connectivity, whereas broadband coherence reflects coordination across different neuronal processes across bands.

The imaginary coherence was calculated as follows. First, we computed the cross-spectrum between the two signals, S_uv_

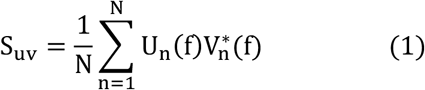

where N refers to the number of samples in the time course, and U and V refer to the Fourier transform of the two signals, respectively.

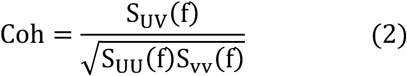

The complex coherence of the signal was calculated as the cross-spectrum normalized by their auto-spectra. Imaginary coherence is the magnitude of the imaginary part of the complex coherence.

#### 2.6.4 EEG complexity index (CI)

The complexity index was calculated as in our previous study (Zhong & Chen, 2022). The order of permutation was set to 2 and the noise threshold was 0.5 relative to the standard deviation of each band, while the scales were chosen as 3, 6, 13, 21 and 62 to represent coarse to fine temporal components of the signal. The sample entropy was used to calculate the entropy for each scale. The complexity index (CI) was calculated using the area under the multiscale entropy curve.

### 2.7 Analysis of associations between fMRI and EEG metrics

The associations between network-averaged rs-fMRI metrics and EEG metrics were assessed using a linear mixed-effects model (LME) approach, both before and after macrovascular correction. As in our previous study (Zhong et al., 2025), sex was also included as a fixed effect, since sex effects have been observed in both rs-fMRI (Tomasi & Volkow, 2012) and EEG metrics (Nentwich et al., 2020; Zhong & Chen, 2022). The z-transform was applied to all parameters prior to model fitting. Outliers were detected and removed for each fitting according to the 1.5 interquartile range (IQR) criterion. An overview of the parameters of interest is presented in **Table 1** and the model is described as **Eq. 3**. A significance level of 0.05 was used with a false discovery rate (FDR) correction for results of pre- and post-macrovascular correction separately. The goodness-of-fit (r^2^) values for the models are presented with the number of coefficients adjusted. The same model was also used for examining associations within each network. Moreover, in order to confirm that the associations from the LME model reflect inter-participant differences rather than inter-ROI differences, a similar model with network ID added as a random variable was also built to determine whether this model will result in a difference in effect size from **Eq. 3**.

**Table 1.** Parameters for linear mixed effect model.

| **Y ~ 1 + X + Sex + X:Sex (Eq. 3)** |  |
| --- | --- |
| Y (fMRI) | X (EEG) |
| RSFA | Coherence (Broadband) |
| Seed-based FC | Coherence (Band-limited) |
| gFCD | Entropy |
| IFCD | fractional power (Band-limited) |
| Entropy | Total power (Broadband) |

**Table 2.** Summary of polarity of the associations across EEG metrics. Red: positive effect; blue: negative effect.

| EEG→ | Delta |  | Theta |  | Alpha |  | Beta |  | Gamma |  | broadband |  |  |
| --- | --- | --- | --- | --- | --- | --- | --- | --- | --- | --- | --- | --- | --- |
| fMRI↓ | Power | Coherence | Power | Coherence | Power | Coherence | Power | Coherence | Power | Coherence | Power | CI | Coherence |
| gFCD | + | - | N/A | - | - | + | N/A | - | - | - | N/A | - | N/A |
| IFCD | N/A | N/A | + | N/A | N/A | + | N/A | - | - | - | N/A | N/A | - |
| Seed-based FC | + | N/A | - | N/A | - | N/A | N/A | - | N/A | - | N/A | N/A | N/A |
| RSFA | N/A | N/A | N/A | N/A | N/A | N/A | N/A | N/A | N/A | N/A | N/A | - | N/A |
| entropy | - | N/A | N/A | N/A | + | - | + | + | + | N/A | - | + | N/A |

## 3 Results

### 3.1 Variations across networks

Network definitions based on the rs-fMRI-based FC values depended on whether macrovascular correction was applied. This resulted in pre-correction and post-correction sets of network ROIs, which were used for extracting pre- and post-macrovascular correction fMRI metrics, respectively. According to **Figure S1-4**, there was no significant difference across networks for any of the EEG and rs-fMRI metrics, likely due to the high inter-participant standard deviation.

### 3.2 Model goodness-of-fit between post- and pre-macrovascular correction

Goodness-of-fit statistics help to assess the overall quality of the model and determine if it adequately fits the data, whereas effect size reflects the strength of the associations between variables in the model. Among all fMRI metrics, gFCD showed the best goodness of fit to the LME model (as measured by r^2^) for EEG metrics, while the seed-based FC and fMRI entropy showed the worst. For band-limited EEG metrics (both fractional power and coherence), the r^2^ varied according to the rs-fMRI metric used, but in the majority of cases, the alpha and beta bands outperformed other bands in terms of r^2^. The only exception was for associations between gamma coherence and rs-fMRI lFCD, which showed a higher r^2^ than all other EEG coherences. Additionally, among broadband EEG metrics, EEG CI was found to fit most rs-fMRI metrics better than other EEG metrics (the only exception being lFCD).

Across all model fits shown in **Figures 2 to 4**, macrovascular correction improved the r^2^ values. The only exceptions pertain to very poor fits (low r^2^). Among the associations with reasonable effect sizes, the percent improvement in r^2^ was highest for lFCD, followed by seed-based FC, with regard to all EEG metrics.

**Figure 1.**
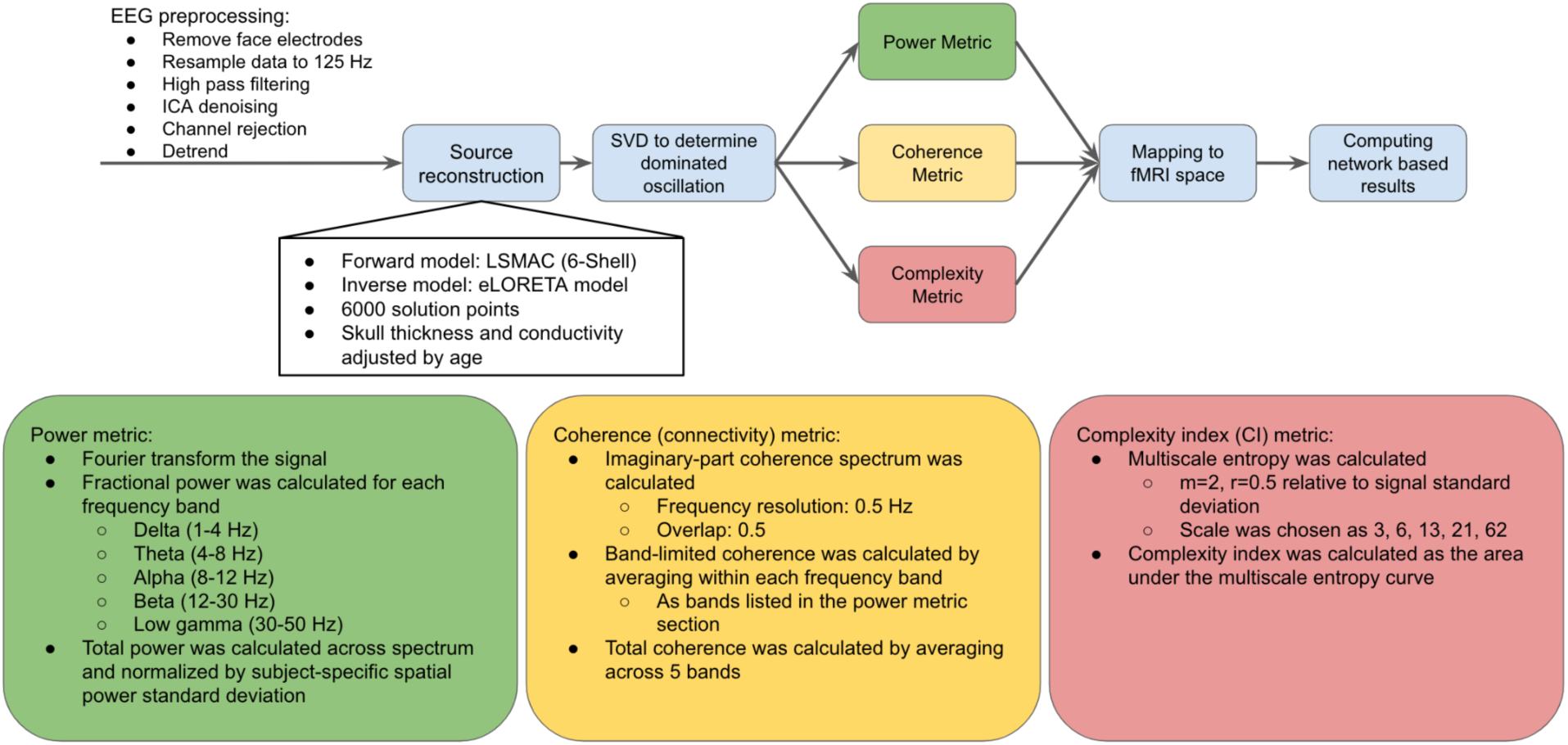
EEG preprocessing and analysis pipeline.

**Figure 2.**
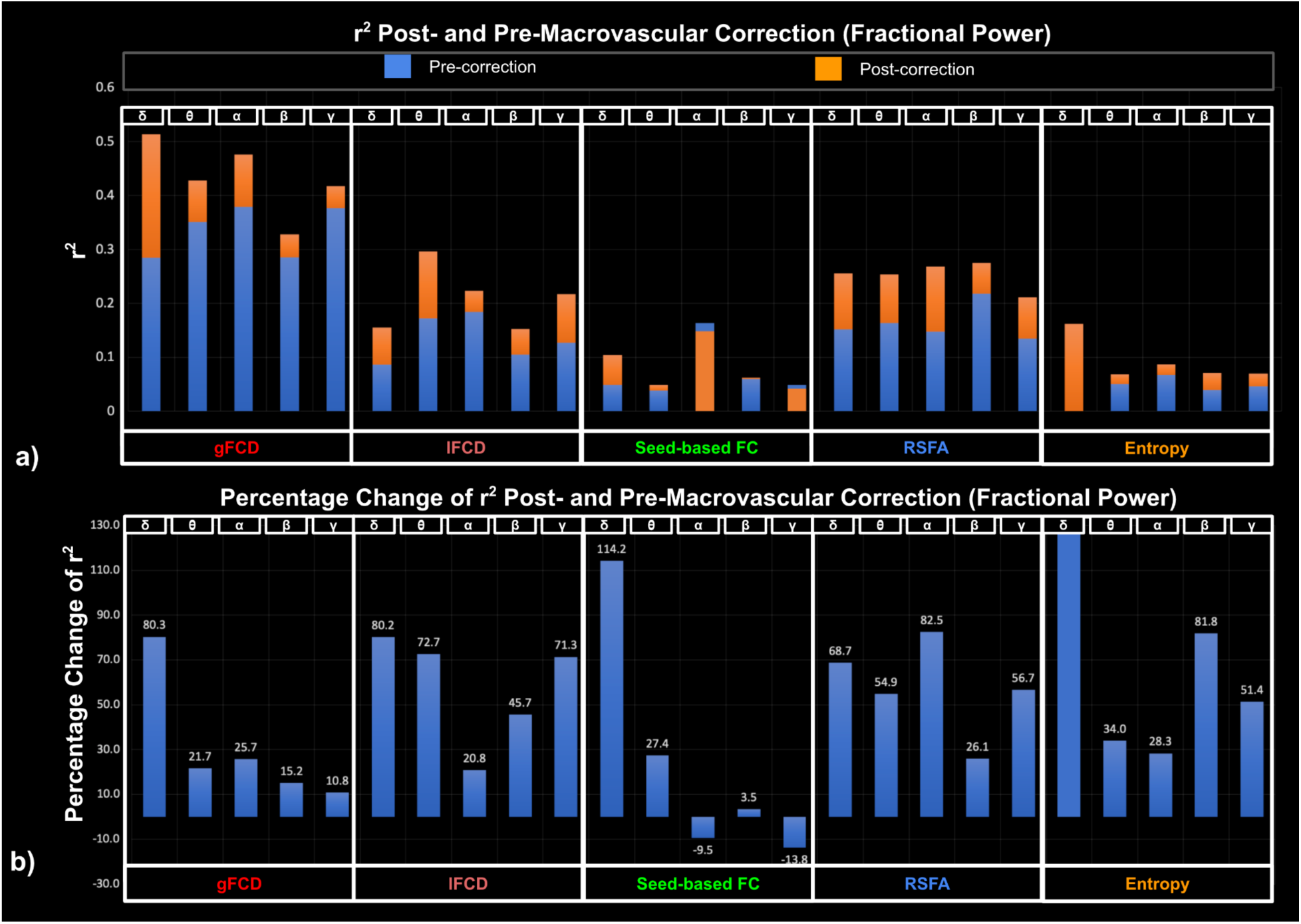
Goodness of fit (r^2^) of LME models for EEG fractional power, compared between post- and pre-macrovascular correction. a) r^2^ post- and pre-macrovascular correction; b) percentage difference in the r^2^ post- and pre-macrovascular correction;

**Figure 3.**
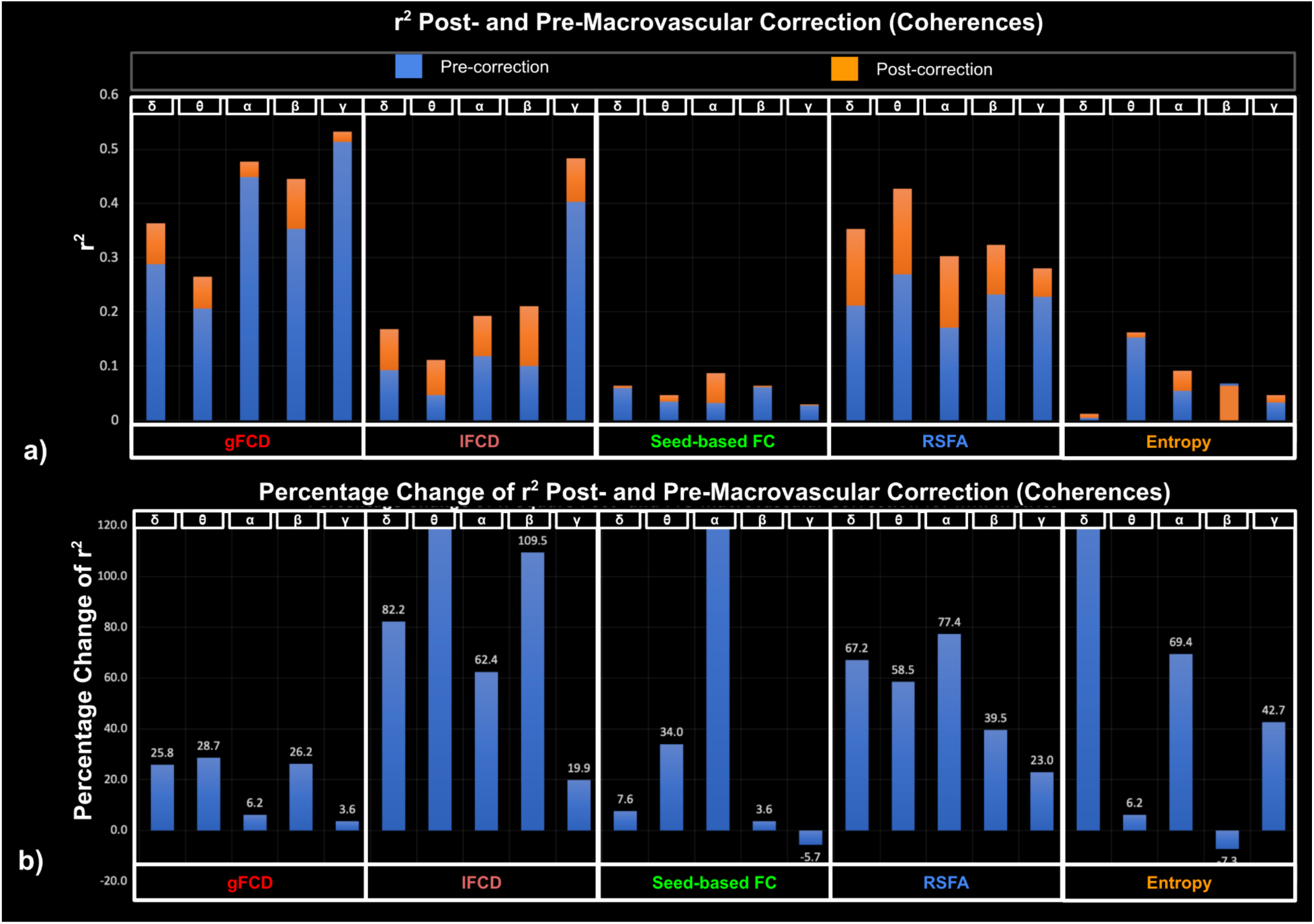
Goodness of fit (r^2^) of LME models for EEG coherence, compared between post- and pre- macrovascular correction. a) r^2^ post- and pre-macrovascular correction; b) percentage difference in the r^2^ post- and pre-macrovascular correction.

**Figure 4.**
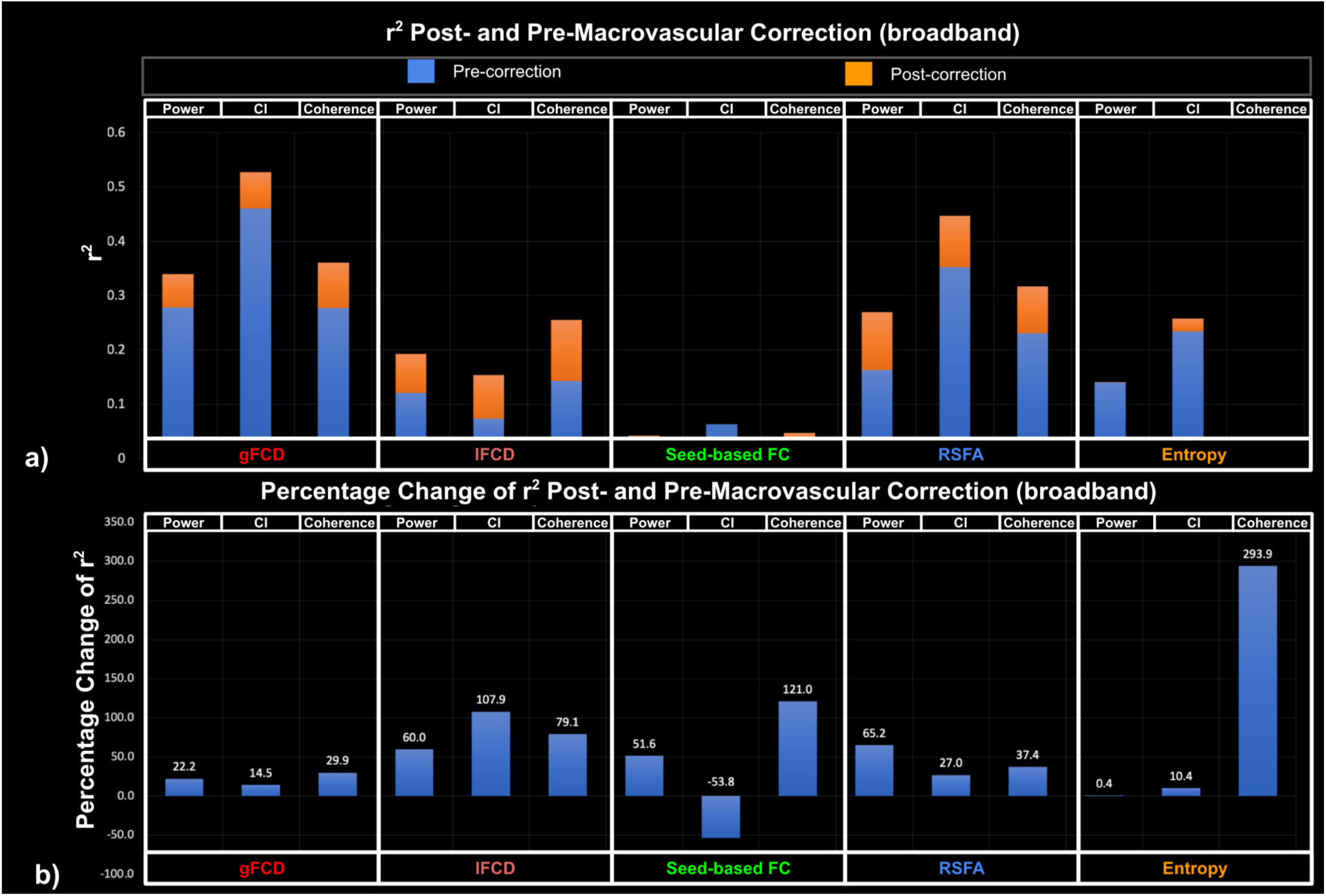
Goodness of fit (r^2^) of LME models for EEG broadband metrics, compared between post- and pre-macrovascular correction. a) r^2^ post- and pre-macrovascular correction; b) percentage difference in the r^2^ post- and pre-macrovascular correction.

While the associations between seed-based FC and rs-fMRI entropy with EEG metrics showed the greatest improvement after macrovascular correction (**Fig. 2-4**), they also showed poor r^2^ to the rs-fMRI FC metrics both pre- and post-correction, despite this strong r^2^ improvement. There was a marked improvement in associations with EEG metrics for rs-fMRI lFCD and RSFA after macrovascular correction, bringing r^2^ up to 0.35. Lastly, the associations between gFCD and EEG metrics, which were already strong pre-correction, were further improved after macrovascular correction.

### 3.3 Association between fMRI and EEG metrics

#### 3.3.1 Comparison of rs-fMRI-EEG associations post and pre-macrovascular correction

We observed negligible differences in effect sizes with and without the network ID as a random variable. Thus, the associations were dominated by inter-participant differences, and are deemed largely consistent for different networks. This is supported by the results for individual networks shown in **Fig S5-7**.

Figures 5-7 illustrate how the significance and size of the rs-fMRI-EEG associations could vary between pre- and post-macrovascular correction. In the case of band-limited coherence, the association between the beta coherence and lFCD was only significant with macrovascular correction. Furthermore, sex differences only became significant after macrovascular correction, as seen in the associations between delta band fractional power and rs-fMRI entropy, as well as between delta coherence and RSFA. (Figure 5). Likewise, the association between the gamma fractional power and lFCD became significant after macrovascular correction, while the association between the alpha fractional power and lFCD did not survive. Moreover, in most cases (delta, alpha and beta bands), associations between EEG band-limited fractional power and fMRI entropy were only significant after macrovascular correction (Figure 6). With regard to broadband EEG metrics, the association between EEG power and fMRI entropy was only significant after macrovascular correction, while the association between EEG power and lFCD did not survive the macrovascular correction (Figure 7).

**Figure 5.**
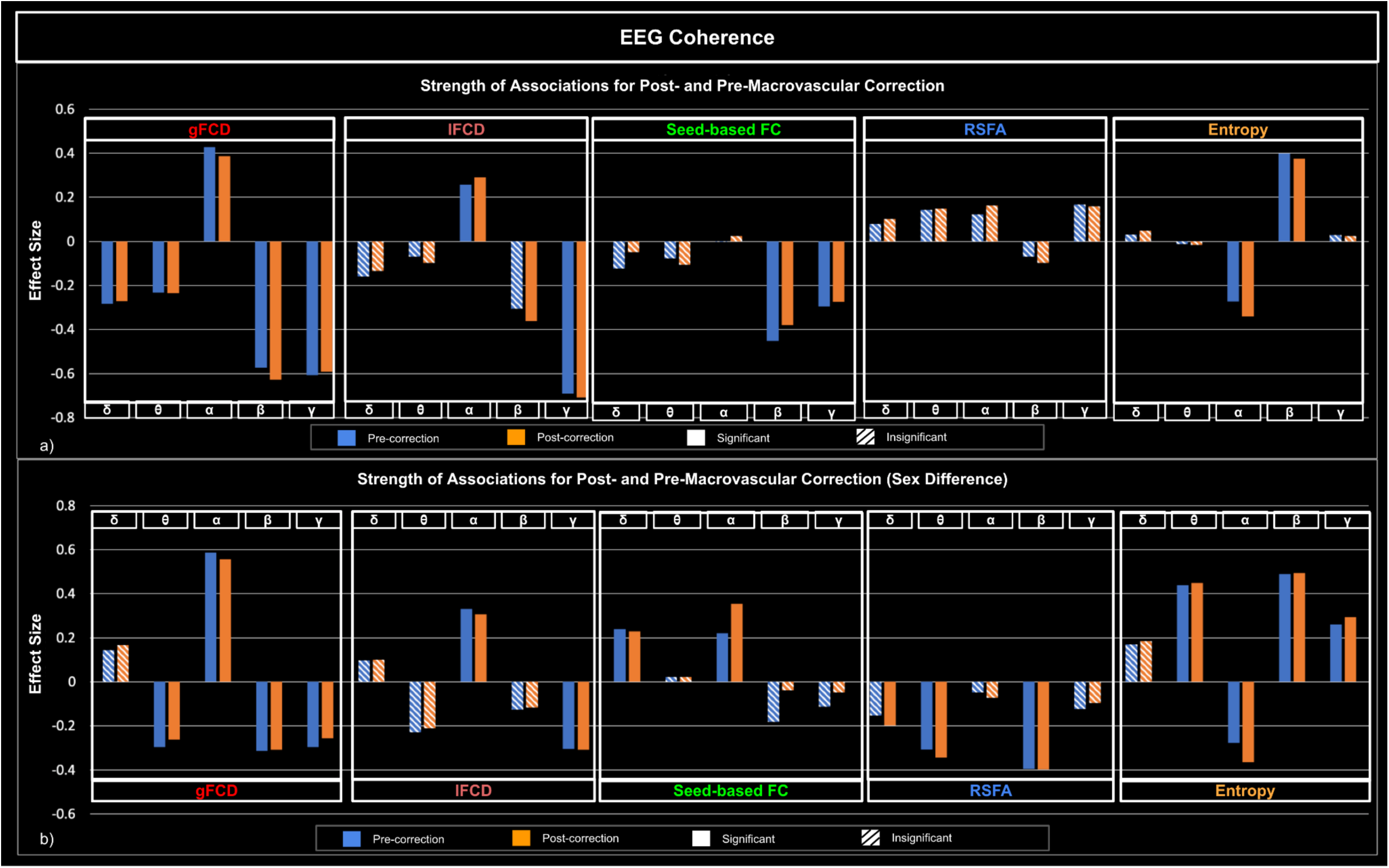
Associations between rs-fMRI metrics and EEG band-limited coherence, with and without macrovascular correction. a) Strength of associations between rs-fMRI metrics and EEG band-limited fractional power; b) comparison of strength of associations between males and females, with a positive effect size indicating stronger associations in males. Blue: pre-correction; orange: post-correction; solid bar: significant association; striped bar: insignificant association.

**Figure 6.**
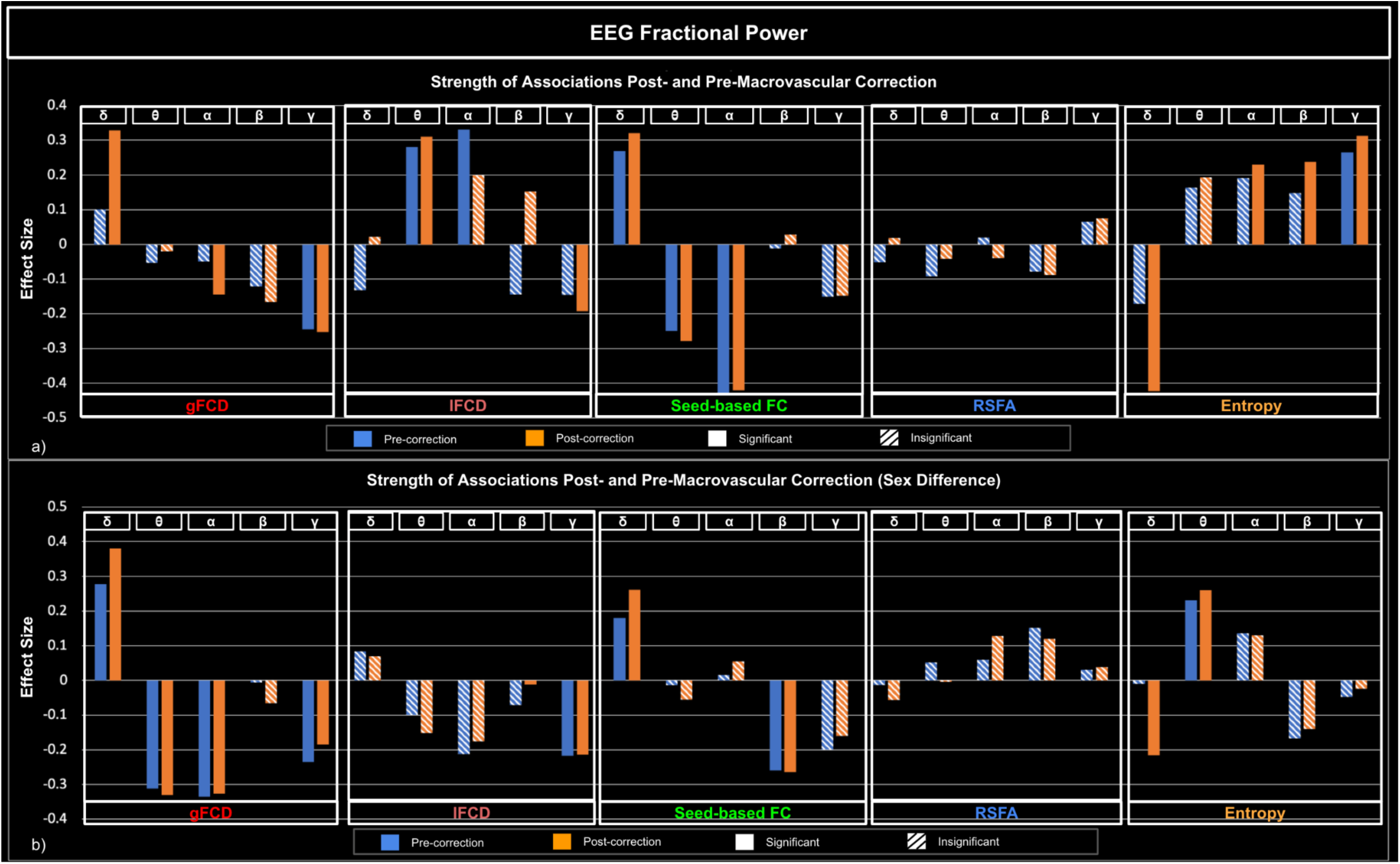
Associations between rs-fMRI metrics and EEG band-limited fractional power, with and without macrovascular correction. a) Strength of associations between rs-fMRI metrics and EEG band-limited fractional power; b) comparison of strength of associations between males and females, with a positive effect size indicating stronger associations in males. Blue: pre-correction; orange: post-correction; solid bar: significant association; striped bar: insignificant association.

**Figure 7.**
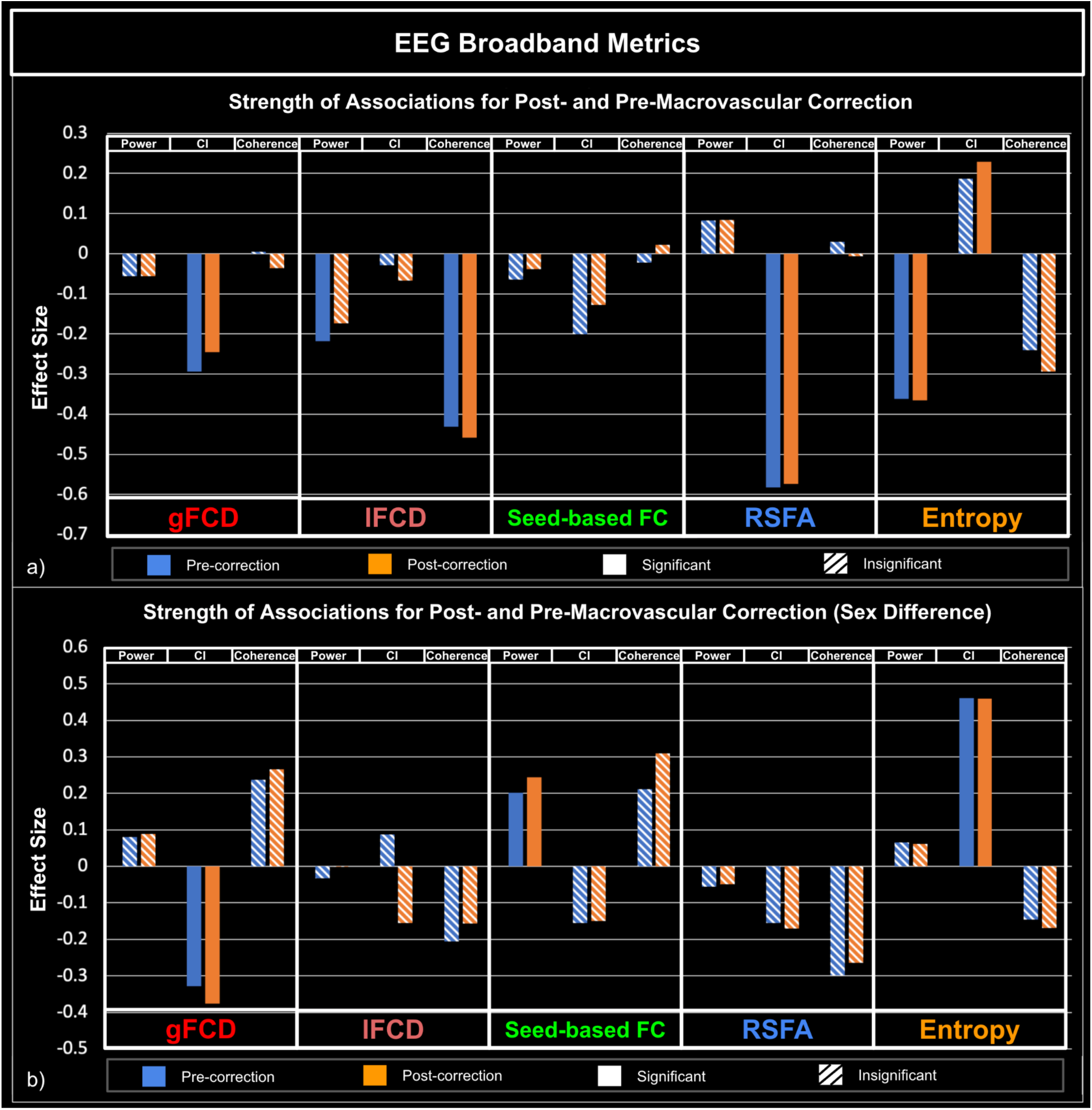
Associations between rs-fMRI metrics and EEG broadband metrics (Total power, entropy and coherence), with and without accounting for macrovascular correction. a): Strength of associations between rs-fMRI metrics and EEG broadband metrics; b): Comparison of strength of associations between males and females, with a positive effect size indicating stronger associations in males. Blue: pre-correction; orange: post-correction; solid bar: significant association; striped bar: insignificant association.

Unlike the goodness-of-fit measures shown in the previous subsection, the effect size assesses the contribution of each variable to the model rather than the degree to which the LME fits the data. The findings in this section are determined across all functional networks.

#### 3.3.2 EEG band-limited metrics

Similar to the case of fractional EEG power, band-limited coherence was strongly associated with fMRI metrics in a band-dependent way (Figure 5a). gFCD was significantly negatively associated with the coherence of the delta, theta, beta and gamma bands positively associated with the coherence of the alpha band. These bi-directional relationships were observed with lFCD, where the association with alpha coherence was significantly positive, whereas the association with beta and gamma coherence was significantly negative. A significant negative association was also found between the coherence of the beta and gamma bands and fMRI seed-based FC. A significant negative relationship was found between alpha coherence and fMRI entropy, whereas a significant positive relationship was found between beta coherence and fMRI entropy. Again, there was no association between EEG band-limited coherences and fMRI RSFA.

As with band-limited fractional power, sex had a significant effect on the association between EEG band-limited coherences and rs-fMRI metrics (Figure 5b). Accordingly, females showed stronger associations between theta/beta/gamma coherence and gFCD, whereas males showed a stronger association between alpha coherence and gFCD. Males also showed a stronger association between alpha coherence and lFCD, while females showed a stronger association between gamma coherence and lFCD. Coherence in the delta and alpha bands was also found to be more strongly associated with seed-based FC in males. In spite of the fact that there was no association between RSFA and band-limited coherence, there was significant sex difference, in that females showed stronger associations than males between RSFA and coherence in delta, theta and beta bands. For fMRI entropy, males showed stronger associations with coherence in the theta, alpha, and gamma bands, while females showed a stronger association with coherence in the alpha band. The breakdown of associations mentioned above in each individual network is presented in **Fig S5**.

There were significant band-dependent associations between EEG fractional power and rs-fMRI metrics. It was found that the fractional power of the alpha and gamma bands was significantly negatively associated with gFCD, whereas the fractional power of the delta band was significantly positively associated (Fig. 6a). Moreover, the fractional power in the theta and gamma was associated with lFCD, which was positive for theta but negative for gamma. The fractional power of the delta band showed a significant positive association with seed-based FC, while the fractional power of the theta and alpha bands showed a significant negative association. EEG fractional power was not significantly associated with RSFA in any band. Furthermore, we found that the fractional power of the delta band was significantly negatively associated with fMRI entropy, whereas the fractional power of the alpha, beta and gamma bands were significantly positively associated with fMRI entropy.

Sex was found to significantly affect the associations between EEG fractional power and rs-fMRI metrics but in a highly diverse way, as can be seen in Figure 6b. First, the association between EEG delta band fractional power and a number of rs-fMRI metrics, including the gFCD, seed-based FC, and fMRI entropy (Figure 6b). Specifically, males have stronger associations with gFCD and seed-based FC, whereas females have stronger associations with fMRI entropy. Similar sex effects were also observed for theta band fractional power (females show a stronger association with gFCD, males show a stronger association with rs-fMRI entropy), alpha band fractional power (females show a stronger association with gFCD), and gamma band fractional power (females show a stronger association with gFCD and lFCD). Interestingly, there was no significant sex effect found in the association between beta band fractional power and any of the rs-fMRI metrics. The breakdown of associations mentioned above in each individual network is presented in **Fig S6**.

#### 3.3.3 EEG broadband metrics

The association between broadband EEG matrics and fMRI metrics was limited (Fig. 7a) and primarily through EEG entropy. There was a significant negative association between the EEG CI and the gFCD as well as RSFA. There was also as a significant positive association between the EEG CI and the fMRI entropy. A significant negative association was also found between EEG coherence and lFCD. Furthermore, EEG power was found to be significantly negatively associated with both fMRI entropy and lFCD.

Similarly, associations in broadband EEG metrics were significantly affected by sex (Figure 7b). In males, EEG CI was more strongly associated with fMRI entropy, and in females, EEG CI was more strongly associated with gFCD. The breakdown of associations in each individual network is presented in **Fig S7**.

## 4 Discussion

How rs-fMRI measurements are related to EEG counterparts has long been considered a key question for understanding the neuronal underpinnings of rs-fMRI measurements. Despite this, the presence of venous bias in rs-fMRI measurements makes it difficult to accurately identify such associations. Recently, we demonstrated a macrovascular correction approach that effectively improved the goodness-of-fit in that model that explains rs-fMRI measurements using metabolism and perfusion (Zhong et al., 2025). In this study, we apply the same macrovascular bias correction in investigating the link between rs-fMRI and EEG measurements. Our main results are:

1. Macrovascular correction improved the goodness-of-fit in the models relating EEG metrics and rs-fMRI metrics. This is in agreement with Hypothesis 1.
2. Sex appears to have a significant impact on the EEG-fMRI associations. This is in line with previous studies and agrees with our Hypothesis 2.
3. The EEG complexity was significantly associated with the RSFA, which agrees with our Hypothesis 3. This association, however, is negative, rather than positive as found in the brain aging context (Zhong & Chen, 2022).

We also examined the associations between various EEG metrics and rs-fMRI metrics, particularly the associations across the different EEG frequency bands as well as in broadband EEG metrics. We found:

### 4.1 Effect of the macrovascular correction

As our recent work shows, macrovascular contributions inflated BOLD signal power and functional connectivity metrics, which can be evident even at > 1 cm away from the vascular boundary at typical fMRI voxel sizes (Zhong, Tong, et al., 2024). These inflated BOLD signal strengths and functional connectivity values are unlikely to be neuronal in origin (Zhong, Tong, et al., 2024). The non-neuronal contribution to the rs-fMRI measurements adds to discrepancies between the EEG and rs-fMRI metrics (assuming that rs-fMRI and EEG reflect the same neuronal activity). Thus, macrovascular BOLD contributions may be one of the major reasons why previous studies have reported inconsistent relationships between EEG and rs-fMRI metrics. In our recent work, we demonstrated that macrovascular correction substantially reduces the bias on the BOLD metrics; specifically, gFCD was reduced globally, while the lFCD and RSFA were reduced locally (Zhong et al., 2025). As a result, we showed that rs-fMRI metrics became better modeled by CMRO_2_. By extension, we hypothesize the same when relating rs-fMRI to EEG metrics.

In this work, we show that the r^2^ relating rs-fMRI metrics and EEG metrics also improved, supporting our hypothesis. A higher r^2^ after macrovascular correction would suggest that rs-fMRI metrics are closer to their EEG counterparts and can better reflect local-specific neural activity (Zhong et al., 2025) - this is exactly what we observed in all cases. Across all fMRI metrics, the EEG-associations were most improved for lFCD and RSFA. This finding echoes findings from our previous study (Zhong et al., 2025), in which we noted that lFCD and RSFA are the most susceptible to macrovascular bias, and correspondingly their associations with CMRO_2_ show stronger improvement than other metrics after macrovascular correction. However, macrovascular correction only brought about limited improvement and in some cases, a reduced r^2^, in the EEG-association for rs-fMRI seed-based FC and entropy metrics. Since the r^2^ values for these two metrics were low initially (i.e. <0.1, Fig. 2**-4**), the extent to which these two metrics can be explained by EEG metrics is unclear. This also echoes associations of these two metrics with CMRO_2_ in our previous study (Zhong et al., 2025). A further point worth mentioning is the fact that gFCD showed the highest r^2^ values along with the lowest r^2^ improvement after macrovascular correction, which could indicate that gFCD would be the metric most immune to macrovascular bias.

In line with our study regarding CMRO_2_ and its correlates (Zhong et al., 2025), applying macrovascular correction altered the significance of the associations between EEG metrics and rs-fMRI metrics. In the example of delta band fractional power and gFCD, the associations only become significant after macrovascular correction, and similarly for the association between gamma band fractional power and lFCD as well as between beta band coherence and lFCD of rs-fMRI. Such increases in significance of the EEG-fMRI associations could provide new insights when investigating the relationship between EEG and rs-fMRI. This is also the case for associations that transition from significant to insignificant after macrovascular correction, which may in some cases suggest false positives in the past due to macrovascular bias in fMRI. Furthermore, in the associations between EEG fractional power and rs-fMRI entropy, numerous associations transitioned from insignificant to significant after macrovascular correction, strongly suggesting that macrovascular correction is necessary while investigating the rs-fMRI entropy, even though the mechanism that contributes to entropy remains unclear. Lastly, unlike in the case of CMRO_2_ (Zhong et al., 2025), the strength of EEG-fMRI associations varied widely across different EEG metrics, as will be discussed in the next section.

### 4.2 Associations between fMRI metrics and band-limited EEG metrics

It is widely accepted that EEG measurements can be divided into different frequency bands, which may correspond to different aspects of brain activity (Bouafif, 2021). Therefore, band-limited approaches are intuitive in establishing links between EEG and rs-fMRI (Scheeringa et al., 2012; Xavier et al., 2025). The EEG power and connectivity have been investigated previously, but with limited understanding of their respective associations to diverse rs-fMRI metrics, as discussed earlier. In this study, we focused on inter-participant associations between static EEG and rs-fMRI metrics, which can provide additional information than previous studies that examined associations across regions and/or time points.

#### 4.4.1 EEG Band-limited coherence vs. rs-fMRI connectivity

Even though rs-fMRI and EEG measure neuronal activity via different mechanisms, it is natural to use the association between connectivity metrics derived from these two methods to understand the underpinnings of rs-fMRI FC. In this case, EEG coherence is taken as a widely accepted measure of EEG-based FC. As Previous work showed that phenotype, particularly the sex effect, fundamentally modulated the inter-region association between EEG coherence and rs-fMRI FC (Nentwich et al., 2020), we included sex as a covariate in determining the EEG-fMRI connectivity associations. We found that EEG coherences across different frequency bands were associated with multiple rs-fMRI metrics across participants, with the most widespread EEG associations found for the rs-fMRI gFCD metric. Moreover, we found that the alpha was the only band with coherence showing positive associations with rs-fMRI FC metrics (gFCD and lFCD), which suggests that alpha connectivity has the strongest association with rs-fMRI FC. This is consistent with previous whole-brain findings using EEG and MEG showing that alpha and beta connectivity (Brookes et al., 2011; de Pasquale et al., 2010; Garcés et al., 2016; Wirsich et al., 2021) are most strongly associated with rs-fMRI FC, albeit these were inter-regional rather than inter-participant associations. Our findings are also consistent with previous interareal findings of alpha connectivity being most strongly associated with rs-fMRI FC in the VN (L. Wang et al., 2012). On the other hand, we did not find strong positive associations between gamma connectivity and rs-fMRI FC. Gamma rhythm, which has previously been associated with rs-fMRI (Magri et al., 2012), is more strongly associated with rs-fMRI FC across shorter distances (Deligianni et al., 2014; Wirsich et al., 2017).

It is interesting to note that with the exception of alpha band, the EEG coherence showed a negative association with rs-fMRI FC. There are several factors potentially contributing to such findings. First, macrovascular correction, which was not done in previous studies, may have an important role in these associations. Secondly, our findings are driven mainly by inter-participant differences in EEG and fMRI measurements, unlike previous studies, which mostly examine inter-regional differences. We will discuss these points in more detail as follows.

First, as we discussed in the “e*ffect of the macrovascular correction”* section, by removing the macrovascular signal, we expect rs-fMRI measurements to more closely reflect their neuronal contributions and thus, EEG measurements. l. Previous findings suggest that fMRI measures of brain activity overlap more with the portion of the EEG that is stationary (de Pasquale et al., 2010). Resting-state macrovascular fMRI signal is likely to be more stationary and global (Zhong, Tong, et al., 2024) than the underlying neuronal activity. This is potentially due to the contribution of systematic low-frequency oscillations (Zhong, Tong, et al., 2024), which correspond to physiological respiratory and cardiovascular components (Tong et al., 2019). By removing the macrovascular contribution, we reduce the stationarity in the fMRI signal and strengthen the contribution from non-stationary fMRI components that could more closely reflect the EEG signal. Further, previous research has indicated that delta and theta connectivity patterns exhibit the greatest spatial similarity with rs-fMRI-based functional networks, since both bands display a greater degree of global (interhemispheric) connectivity (Deligianni et al., 2014; Wirsich et al., 2017). However, as we have shown, macrovascular bias contributes to a substantial portion of the global connectivity of rs-fMRI (Zhong, Tong, et al., 2024), which has previously led to a positive association between rs-fMRI FC and low-frequency EEG FC. However, there remain major methodological differences between our study and those reporting positive EEG-fMRI FC associations, and the effect of macrovascular contributions above and beyond these differences requires further investigation.

Secondly, the EEG-fMRI associations may differ when driven by inter-participant versus inter-regional differences. A systematic difference in delta FC across individuals, potentially driven by differences in brain states or levels of cognitive functioning across participants (e.g., different attention levels) (Clarke et al., 2007), may alter the EEG-fMRI association relative to within-participant inter-regional association. Specifically, reduced delta coherence (Clarke et al., 2007) in the presence of increased rs-fMRI FC (Zhang et al., 2024) has been found for ADHD patients. Moreover, in the process of brain aging, EEG beta coherence (Vysata et al., 2014) increases while rs-fMRI global FC declines (Xie et al., 2020), which results in a negative association between the two modalities. Thus, our findings support the idea that the associations between the two metrics among healthy young adults could be a more moderate version of those seen in healthy aging or between populations.

A further interesting finding was that the associations between each band depended on which connectivity metrics were used in rs-fMRI and, more specifically, what spatial scale was used in rs-fMRI FC. As a result of the biophysical properties of electrical signals, longer conduction delays between distant brain regions may limit the frequency of long-distance network oscillations to below gamma because high-frequency signals are easier to attenuate during transmission (Miller, 2007; Siegel et al., 2012). Another way to put it is that the slow bands (i.e. delta and theta bands) should be more prominent in long-distance communications (Lu et al., 2007; Pan et al., 2011) while higher-frequency bands such as gamma should be more prominent in short-distance communication. However, when the spatial scale between the two extreme example listed above (for example, network connectivity), the alpha (L. Wang et al., 2012) or the beta (Wirsich et al., 2021) band may be responsible for the connectivity. We have indeed found the alpha band to be the only positive association between gFCD and lFCD (insignificant for seed-based FC), suggesting both gFCD and lFCD may both reflect alpha-like medium-range connectivity.

#### 4.4.2 EEG band-limited fractional power vs. rs-fMRI connectivity

EEG fractional power represents the activity amplitude in each frequency band relative to total activity across all EEG bands. As mentioned earlier, we used fractional EEG power to neutralize the effect of total power fluctuations. This approach is particularly suited to the determination of EEG-fMRI association while the two sets of data were not acquired simultaneously. EEG power is generated by synchronous neuronal activity, and when neuronal activity across brain regions is synchronized, it can be reflected in higher EEG FC. Thus, it is reasonable to expect EEG power to be associated with EEG coherence, and by extension, rs-fMRI FC, although EEG power and EEG/fMRI FC are local and non-local indices of synchrony, respectively. This has been the rationale in a number of previous studies which associated EEG power with fMRI FC across time points (Scheeringa et al., 2012; Xavier et al., 2025).

Our results showed that the EEG band-limited fractional power measurements are significantly associated with multiple rs-fMRI FC metrics, which is consistent with these previous studies. This is despite differences in our methodology, as our results mainly reflect variations in static power across participants instead of power variations across time points. In addition, the associations between EEG fractional power and rs-fMRI FC metrics (gFCD, lFCD and seed-based FC) are bidirectional (i.e. either positive or negative). Although fractional EEG power is unlike absolute EEG power, in that it only represents the contribution of different EEG bands to total power irrespective of changes in total power, our findings are consistent with previous findings based on absolute EEG power across time points (Xavier et al., 2025). This likely indicates that variations in fractional EEG power are consistent with variations in total EEG power across individuals. This consistency between intra-participant and inter-participant associations could suggest that fractional power is less affected by systemic differences in brain activity among individuals. Furthermore, it may also suggest that associations between EEG power and fMRI FC across participants may be based on a similar mechanism as the associations across time points, rather than associations across brain regions.

We note that the associations with three of the rs-fMRI FC metrics are driven by different EEG bands. Specifically, delta activity, besides being associated with sleeping (Purves et al., 2001), has also been linked with cognitive function and has the potential to modulate attention networks. Delta fractional power was most strongly associated with gFCD, whereas theta fractional power was most strongly associated with lFCD. We also noticed that delta fractional power was the only power metric that showed a significant positive association with rs-fMRI FC metrics, especially with gFCD and seed-based FC. This finding is also consistent with data using positron emission tomography (PET), which showed that absolute delta power correlates positively with glucose metabolism in the medial frontal cortex (Alper et al., 2006) gFCD (Shokri-Kojori et al., 2019; Tomasi et al., 2013). This pattern is somewhat consistent with the expectation that both delta band and gFCD encompass more long-range brain-signal contributions, in contrast with lFCD and theta band, which reflect less global activity (see more discussion in *EEG Band-limited coherence vs. rs-fMRI connectivity* section). However, the relationship between fractional EEG power and FC metrics requires more systematic investigation.

We also found a negative association of alpha fractional power with gFCD and seed-based FC. Given that alpha EEG coherence was found to be positively correlated with rs-fMRI FC, the finding regarding EEG fractional power is consistent with previous findings that alpha EEG power is inversely correlated with rs-fMRI FC in multiple brain networks (Chang et al., 2013; Scheeringa et al., 2012). This was suggested to reflect the unique relationship between alpha power and vigilance, such that increased alpha results in higher neuronal inhibition across networks (Chang et al., 2013; Scheeringa et al., 2012). This also suggests that in our results, the inter-participant differences in alpha fractional power are dominated by occipital alpha.

According to our results, the low-gamma fractional power is negatively associated with all three metrics of rs-fMRI FC (i.e. lFCD, gFCD, seed-based FC). These negative associations are inconsistent with previous studies that indicated a positive association between low-gamma absolute power fluctuations dynamic rs-fMRI FC across time points (Tagliazucchi et al., 2012). As this previous study was based on absolute power rather than fractional power, an increase in absolute power may not necessarily translate into an increase in fractional power if other power levels in other bands change proportionately. Specifically, an increase in fractional gamma power may be associated with a decrease in low-frequency power, leading to a negative association between gamma fractional power and rs-fMRI FC (Deligianni et al., 2014). This is particularly true since the gamma band affects fewer regions than lower frequency bands (Tagliazucchi et al., 2012). Moreover, only a few electrodes were included in this previous study to establish the relationship between gamma power and rs-fMRI FC, which may not fully reflect gamma activity. Gamma oscillations have been linked to cortical inhibition in previous studies (Muthukumaraswamy et al., 2009). Therefore, an increase in gamma activity across multiple brain regions might then result in a lower rs-fMRI FC due to widespread neuronal inhibition throughout the brain.

#### 4.4.3 EEG band-limited fractional power and coherence vs. fMRI entropy and RSFA

rs-fMRI entropy has recently been associated with cognitive function (Z. Wang, 2021) and brain aging (Z. Wang & Alzheimer’s Disease Neuroimaging Initiative, 2020). It has been suggested that EEG entropy is associated with rs-fMRI network dynamics (M. Liu et al., 2019; D. J. J. Wang et al., 2018). In light of the similar neuronal basis of EEG and rs-fMRI signals, it would be reasonable to predict that such an association should also exist between rs-fMRI entropy and EEG network dynamics. As in our previous study on an aging population (Zhong & Chen, 2022), no inter-participant association between EEG power and RSFA was found. In contrast, in the current work we found that rs-fMRI entropy was associated with EEG fractional power (in delta, alpha, beta and gamma bands), mirroring the mediation effects observed between EEG entropy and rs-fMRI power across the participants in our previous work (Zhong & Chen, 2022). Although the mechanisms behind these associations between nonlinear and linear metrics are still unclear, such findings, in particular, the positive association between fMRI entropy and the fractional power of traditionally cognitive bands, i.e. beta (Lundqvist et al., 2011) and gamma (Fitzgibbon et al., 2004) bands, further support the existence of links between rs-fMRI entropy and neuronal activity (Z. Wang, 2021; Zhong et al., 2025). However, the consistency of these associations across conditions remains unclear.

We observed an association between rs-fMRI entropy and EEG coherence, much as we did with fractional power. Specifically, the positive association between beta coherence and rs-fMRI entropy echoes our finding of similar associations between beta fractional power and rs-fMRI entropy. In light of the fact that beta band is a traditionally cognitive band (Lundqvist et al., 2011), the association between beta coherence and entropy further supports a cognitive explanation for the variation in rs-fMRI entropy. It is interesting to observe a negative association between alpha coherence and rs-fMRI entropy. This may be due to our use of naturalistic resting-state, which may increase occipital alpha coherence (Jaeger et al., 2023) while decreasing rs-fMRI entropy (Camargo et al., 2023). The mechanism behind the association between EEG coherence and rs-fMRI entropy is also unknown. The interpretation and utility of rs-fMRI entropy require further investigation.

### EEG broadband metrics vs. rs-fMRI metrics

Although EEG analysis has conventionally been band-specific, broadband metrics capture overlapping neural oscillations that might be missed when focusing on individual bands in isolation, as suggested by previous studies (Wirsich, Giraud, et al., 2020). According to previous findings with local field potentials (LFPs), variations in broadband power reflect the modulation of variance in the LFP time series, whereas variations in band-limited power reflect the modulation of oscillatory activity (Manning et al., 2009). Additionally, from an adenosine triphosphate (ATP)-centric metabolic perspective (Attwell & Laughlin, 2001), total energy consumption and its hemodynamic correlates would be closer to total spontaneous activity rather than individual band activity. Thus, broadband EEG associations with rs-fMRI metrics may offer additional physiological insight. Moreover, broadband EEG signal metrics (coherence and power) have been used as biomarkers in multiple previous studies (Kitzbichler et al., 2009; Nentwich et al., 2020). In particular, oscillatory correlates of spatial and verbal memory processes are not necessarily restricted within one frequency band, but rather as a broadband effect (Ekstrom et al., 2007; Manning et al., 2009). Nonetheless, the understanding of the associations between rs-fMRI metrics and broadband EEG metrics remains limited.

Interestingly, as in our previous study in an aging population (Zhong & Chen, 2022), we found that EEG entropy to be significantly associated with RSFA. According to our previous study, decreasing EEG entropy can mediate RSFA decreases with advancing age, which suggests a positive association between EEG entropy and RSFA. The phenomenon which appears to be a result of a different mechanism since we found a negative association between RSFA and EEG entropy here. The complex nature of the RSFA may be the reason for this. A significant portion of the RSFA could be accounted for by non-neuronal contributions (Tsvetanov et al., 2020). In light of the fact that the EEG entropy also displayed a positive association with the rs-fMRI entropy, it is possible to interpret this in the context of the cognitive reserve effect (Z. Wang & Alzheimer’s Disease Neuroimaging Initiative, 2020); that is, higher EEG entropy represents higher cognitive reserve, which might further transform into lower RSFA (Lin et al., 2021). Thus, the differences between the EEG-fMRI-cognition associations in younger and older individuals may reflect a paradigm shift in how cognition is sustained in aging, and by extension, in disease. Nonetheless, finding an association between EEG entropy and RSFA under a circumstance other than brain aging confirms that RSFA may reflect both physiological and neuronal contributions, even though there was no correlation between RSFA and other EEG metrics (Kumral et al., 2020; Tsvetanov et al., 2015; Zhong & Chen, 2022).

In the same way as the linear-to-nonlinear relationship between the EEG entropy and the rs-fMRI RSFA, the linear-to-nonlinear relationship can also be seen by associating the EEG power with the rs-fMRI entropy. Despite the lack of understanding as to the mechanism, such associations strongly suggest that the relationship between EEG and rs-fMRI may not follow a direct one-to-one relationship (power-to-power or frequency-to-frequency) but instead an indirect relationship (e.g. through entropy). It is possible that these findings could be useful in the future for gaining a better understanding of neuronal mechanisms under rs-fMRI measurements.

We do not have a clear explanation for the negative association between lFCD and EEG broadband coherence. An explanation for this negative association could be that different broadband coherences represent different brain states and cognitive functions among participants. Even though it is not known what the driving factors of broadband coherence variations are, previous findings indicate that broadband coherence is more sensitive to phenotypic differences (sex, age, IQ) than band-limited coherence (Nentwich et al., 2020), leading to inconsistencies in the polarity of this association.

### 4.3 What do we learn from associating resting-state EEG with fMRI

It is a common practice of using EEG as a benchmark to interpret the neuronal underpinnings of rs-fMRI measurements (Bonmassar et al., 1999; Menon et al., 1997; Nentwich et al., 2020; Wirsich, Amico, et al., 2020). Neuronal activity measured by EEG will require energy supplied through metabolism, which will result in metabolic and hemodynamic variations that can be measured by fMRI (Buxton et al., 2004). Therefore, there should be a link between rs-fMRI and EEG measurements if both measurements originate from the same neuronal origin. In this study, we found significant associations between various pairs of rs-fMRI and EEG measurements, indicating that the majority of rs-fMRI and EEG measurements reflect the same neuronal origins. Meanwhile, we also observed discrepancies between rs-fMRI and EEG measurements, with even some negative associations between the rs-fMRI and EEG FC across participants. Aside from the possibility that rs-fMRI signals contain significant amounts of non-neuronal physiological contributions (Tong et al., 2011), is it possible that these discrepancies are linked to neuronal origins that may not be visible for either rs-fMRI or EEG?

Despite the fact that EEG and rs-fMRI are both believed to measure neuronal activity (Sotero & Trujillo-Barreto, 2008), the biophysical underpinnings of the measurements differ substantially. EEG measurements are based on the summation of postsynaptic potentials (St. Louis et al., 2016), and the current generated by these potentials must be directed toward the electrodes of the EEG (Malmivuo et al., 1997). Therefore, any neuronal activity unrelated to synchronized postsynaptic activity would be undetectable from the electrode signals recorded by the EEG. As mentioned above, the relationship between EEG and fMRI is mainly mediated by metabolism. ATP production is ∼6 times more efficient when involving oxidative metabolism, which is closely associated with the hemodynamic response. Based on a rodent experiment, Attwel et al. reported that the activity of the postsynaptic receptor is only associated with 34% of the total grey matter ATP metabolism, while 47% of which will associate with action potentials and 13% with resting potentials (Attwell & Laughlin, 2001). Although the data for human brain metabolic budgets is limited, based on different axons and dendrite lengths as well as synapse-neuron ratios between rodents and humans, the percentage of ATP consumption for postsynaptic activity could be increased to 74% (Attwell & Laughlin, 2001). Therefore, rs-fMRI can also reflect more than just synchronized postsynaptic activity, but any neuronal activity that triggers hemodynamic response through metabolic process. Moreover, there is EEG-invisible activity that can also contribute significantly to fMRI signals by modulating hemodynamic responses through metabolic byproducts (for example, nitric oxide (NO)) (Tu et al., 2024), contributing to the discrepancy between rs-fMRI and EEG measurements.

Conversely, rs-fMRI does not reflect 100% of brain grey matter ATP metabolism either. It has been demonstrated that BOLD signals are tightly linked with CMRO_2_ and CBF changes, with limited correlation with glucose metabolism (CMR_glu_) (Cholet et al., 1997). However, glucose is essential for ATP production, whereas oxygen may not always be metabolized. Aerobic glycolysis, for example, enables the brain to produce ATP without oxygen metabolism even when sufficient oxygen is available (healthy brain) (Vaishnavi et al., 2010). Neuronal activity powered by aerobic glycolysis may not elicit a significant fMRI signal, but it might still be recorded by EEG. This is particularly true for high-frequency neuronal activity, which might be mainly powered by aerobic glycolysis (Theriault et al., 2023). This may explain the lack of positive association between EEG gamma and beta FC and BOLD FC in this study (**Table 1**). Furthermore, rs-fMRI inherently has a low temporal resolution due to the long hemodynamic response time (Glover, 2011), making it difficult to identify components of fast neuronal dynamics in EEG.

Therefore, even if both EEG and rs-fMRI reflect neuronal sources, the discrepancy between measurements of the two modalities will still exist. Even though the consistent results between two modalities provide important information about neuronal underpinnings of rs-fMRI, the mismatch between rs-fMRI and EEG is not simply an indication of non-neuronal contribution in rs-fMRI but rather a possibility of contribution from EEG-invisible neuronal activity. Furthermore, this possibility suggests that the association between EEG and rs-fMRI may not be a direct one-to-one association, but an indirect systematic association. In order to gain a deeper understanding of this systematic EEG-fMRI association, it would be necessary to link both to metabolic measurements.

### 4.4 Limitations

Limitations directly related to the macrovascular correction, mainly involving limitations in TOF imaging acquisition and marco-VAN model simulation, have already been addressed in our previous paper (Zhong et al., 2025; Zhong, Polimeni, et al., 2024; Zhong, Tong, et al., 2024) and will not be repeated here.

A limitation of this study was that rs-fMRI and EEG were not recorded simultaneously. Consequently, we needed to assume that the static properties of neuronal activities were not significantly different between rs-fMRI and EEG scan sessions for each participant. For this reason, we used naturalistic resting state (with synchronized naturalistic video-watching across EEG and fMRI acquisitions) in this study in order to ensure that participants were in a similar mental state between the EEG session and the rs-fMRI session. Moreover, comparisons between non-simultaneous EEG and fMRI recordings have been widely used in many previous studies (Nentwich et al., 2020; Scrivener, 2021). In fact, our results, which are independent of simultaneous recording, can be widely used to link existing and future rs-fMRI findings without the availability of simultaneous EEG.

Moreover, in our model, we included sex to account for inter-participant variabilities in the EEG-fMRI associations. While this likely led to an improved understanding of these associations, previous studies suggested that other factors (e.g. IQ) may affect the associations as well (Nentwich et al., 2020). The association of EEG and fMRI metrics as a function of phenotype enriches our understanding of neural-hemodynamic associations beyond the simple assumption of invariability, and will be part of our future work.

## 5 Conclusion

Associations between rs-fMRI measurements and their EEG counterparts are considered the gold standard for understanding neuronal contributions to rs-fMRI. The findings of previous studies based on inter-regional associations, however, have been limited and inconsistent. Compared to previous findings based on inter-regional associations, the inter-participant association here could provide a more generalizable link between EEG and rs-fMRI. Therefore, these associations could establish a healthy-adult reference for future studies of brain abnormality. Furthermore, the macrovascular correction can reduce the discrepancy between rs-fMRI and EEG measurements, enabling us to better uncover more consistent associations. Based on these improvements, we establish associations between rs-fMRI and EEG across different metrics. It is hoped that a systematic analysis of associations between rs-fMRI and EEG metrics may facilitate its use in clinical research in the future.

## Acknowledgement

The authors would like to acknowledge financial support from Canadian Institutes of Health Research and the Canada Research Chairs Program (JJC) and funding support from Ydessa Hendeles Graduate Scholarship (XZZ). Special thanks to Dr. Danny J. J. Wang and his collaborators for providing DE-pCASL sequence.

## Ethics

The study was approved by Baycrest REB (#20-28).

## Author contributions

Xiaole (Zachary) Zhong: Conceptualization, data curation, methodology, software, investigation, writing-original draft preparation. Hannah Van Lankveld: Data curation, writing-reviewing and editing. J. Jean Chen: Conceptualization, data curation, methodology, software, investigation, writing-original draft preparation, supervision, writing-reviewing and editing.

## Data and Code Availability

The data and code will be made available upon request.

