## Supplemental Figures for "The link between steady-state EEG and rs-fMRI metrics in healthy young adults: the effect of macrovascular correction"

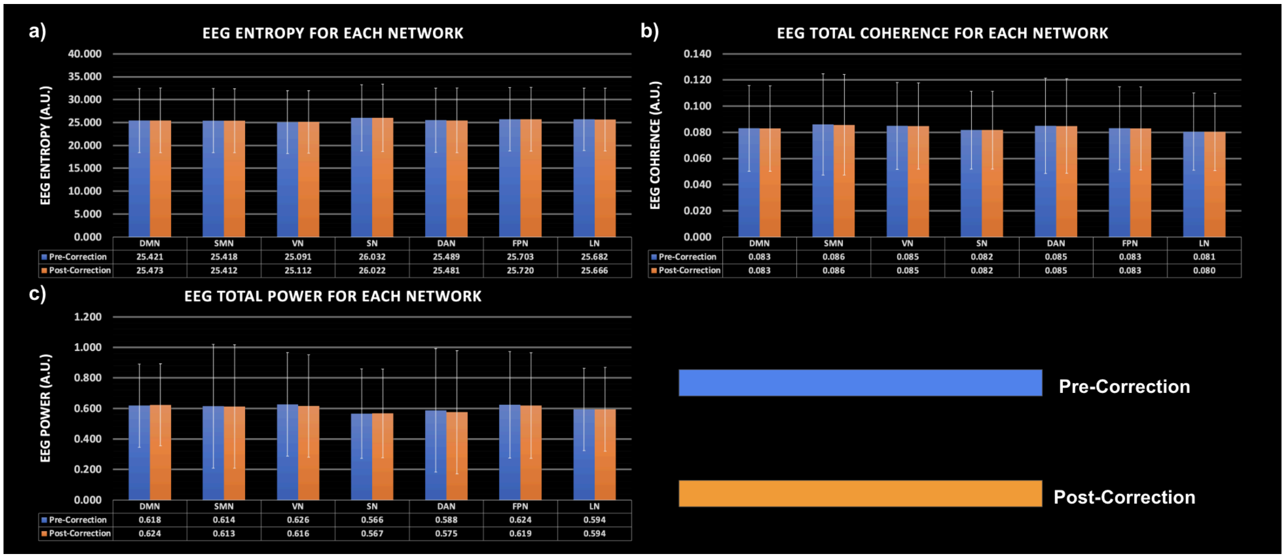


**Figure S1. The average EEG band-free metrics using network ROIs are defined based on rs-fMRI data pre- and post-macrovascular correction.** The EEG metrics corresponding to the post-correction ROIs are of course, corrected for macrovascular contributions. Grouped bars from left to right: DMN, SMN, VN, SN, DAN, FPN, and LN. Blue: pre-macrovascular correction; orange: post-macrovascular correction. Error bars represent the standard deviation.

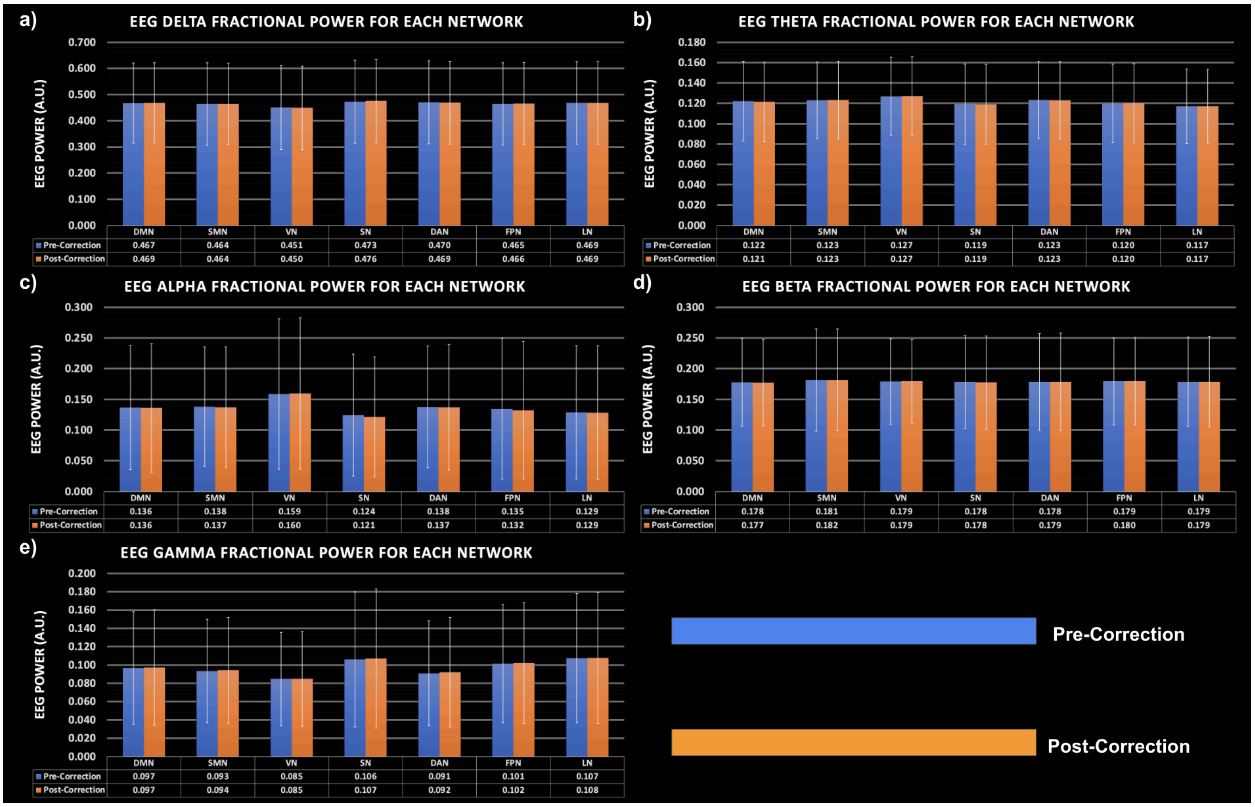


**Figure S2. The average EEG band-limited fractional power using network ROIs is defined based on rs-fMRI data pre- and post-macrovascular correction.** The EEG metrics corresponding to the post-correction ROIs are of course, corrected for macrovascular contributions. Grouped bars from left to right: DMN, SMN, VN, SN, DAN, FPN, and LN. Blue: pre-macrovascular correction; orange: post-macrovascular correction. Error bars represent standard deviation.

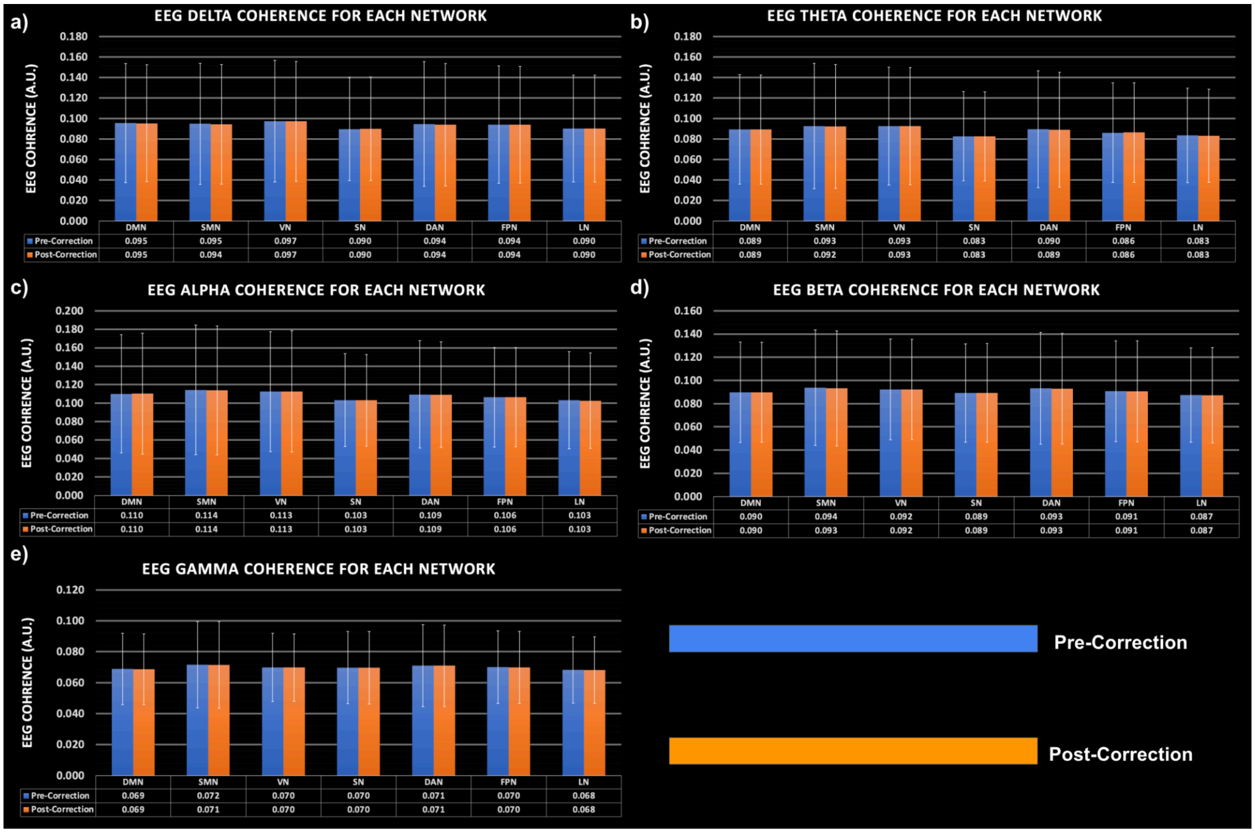


**Figure S3. The average EEG band-limited connectivity (coherence) from network ROIs, computed based on rs-fMRI data pre- and post-macrovascular correction.** The EEG metrics corresponding to the post-correction ROIs are of course, corrected for macrovascular contributions. Grouped bars from left to right: DMN, SMN, VN, SN, DAN, FPN, and LN. Blue: pre-macrovascular correction; orange: post-macrovascular correction. Error bars represent standard deviation.

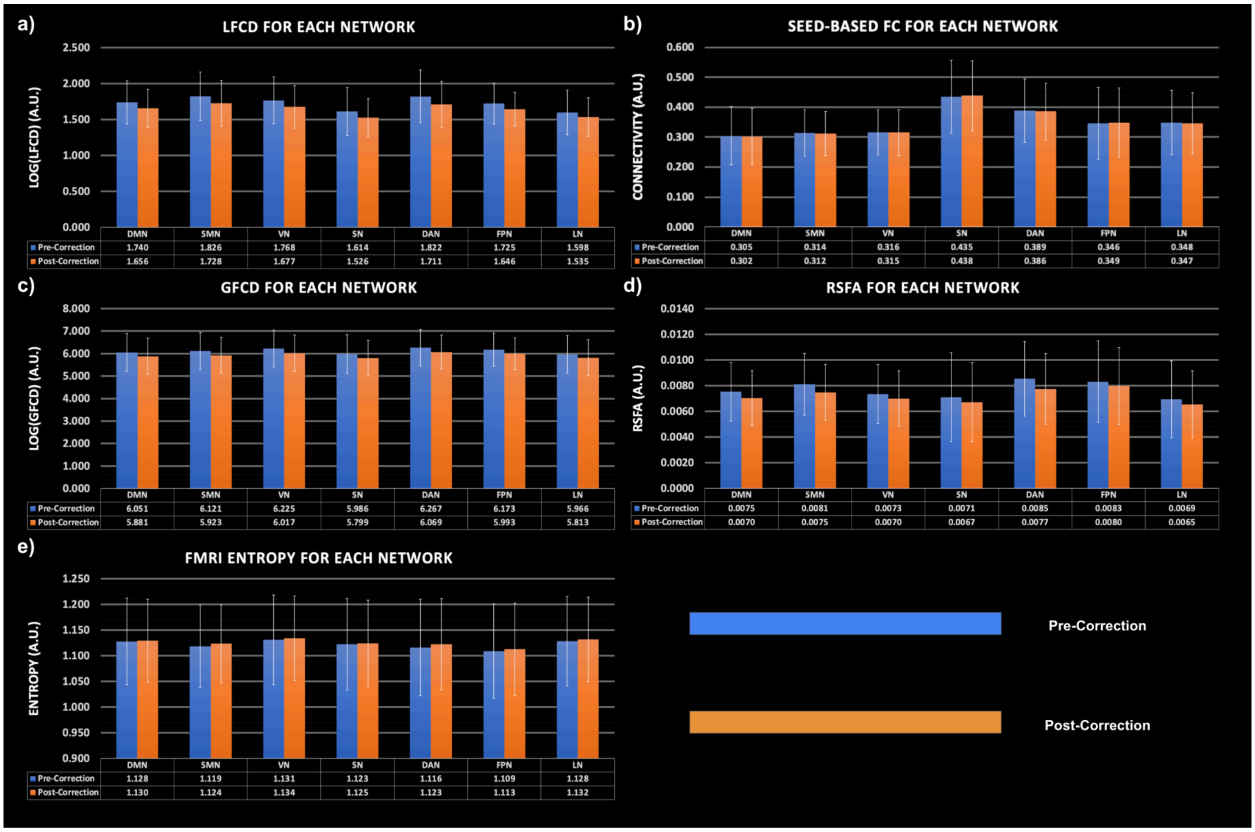


**Figure S4. The average rs-fMRI metrics using network ROIs are defined based on rs-fMRI data pre- and post-macrovascular correction.** The rs-fMRI metrics corresponding to the post-correction ROIs are of course corrected for macrovascular contributions. Grouped bars from left to right: DMN, SMN, VN, SN, DAN, FPN, and LN. Blue: pre-macrovascular correction and orange: post-macrovascular correction. Error bars represent standard deviation.

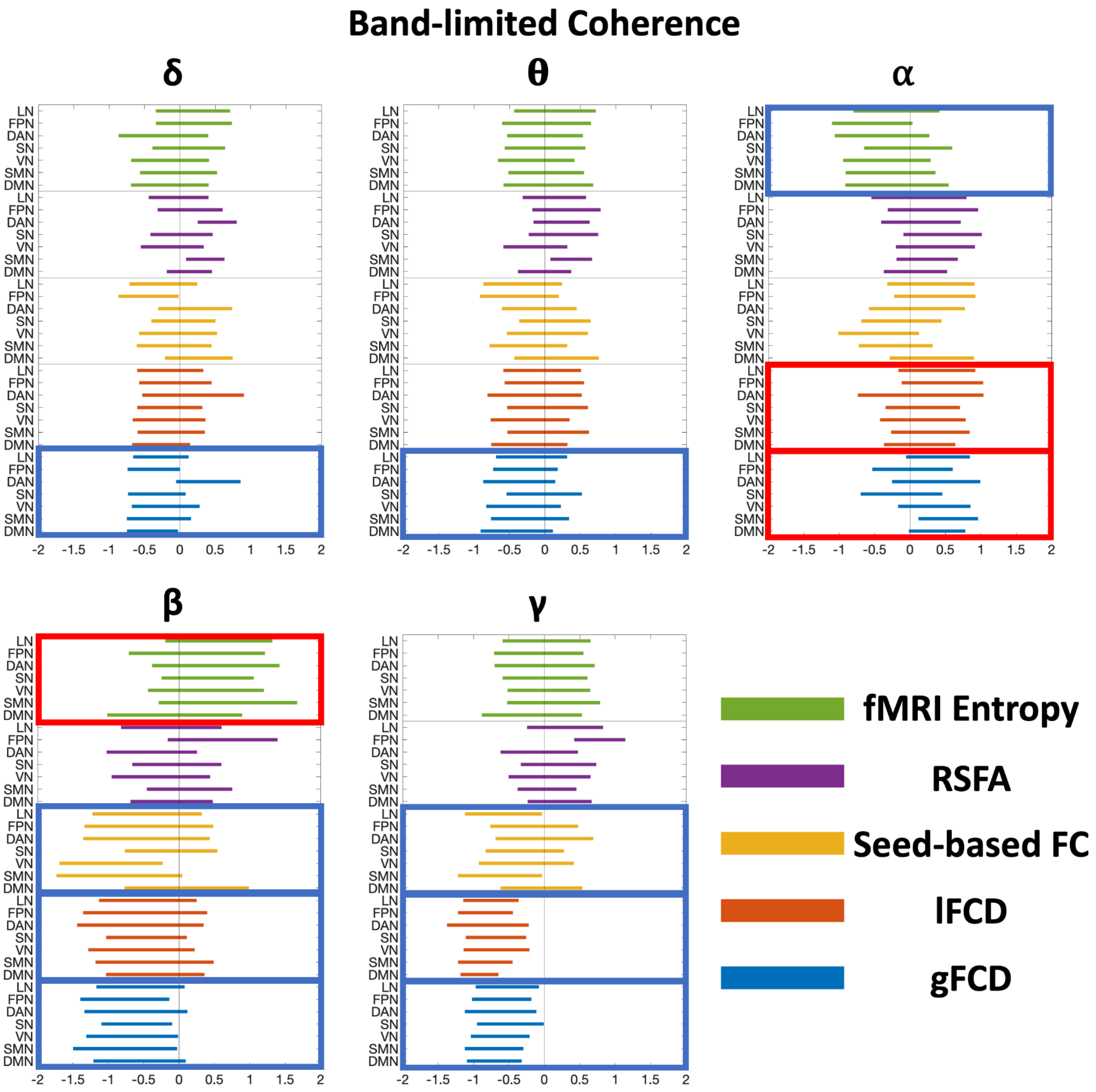


**Figure S5. Effect size of associations between rs-fMRI metrics and EEG band-limited coherence in each individual network, with macrovascular correction.** Each bar represents a 95% confidence interval of the effect size, and its color corresponds to the rs-fMRI metrics as specified by the legend. The box indicates significant whole-brain associations with the color indicating the direction: red for positive and blue for negative.

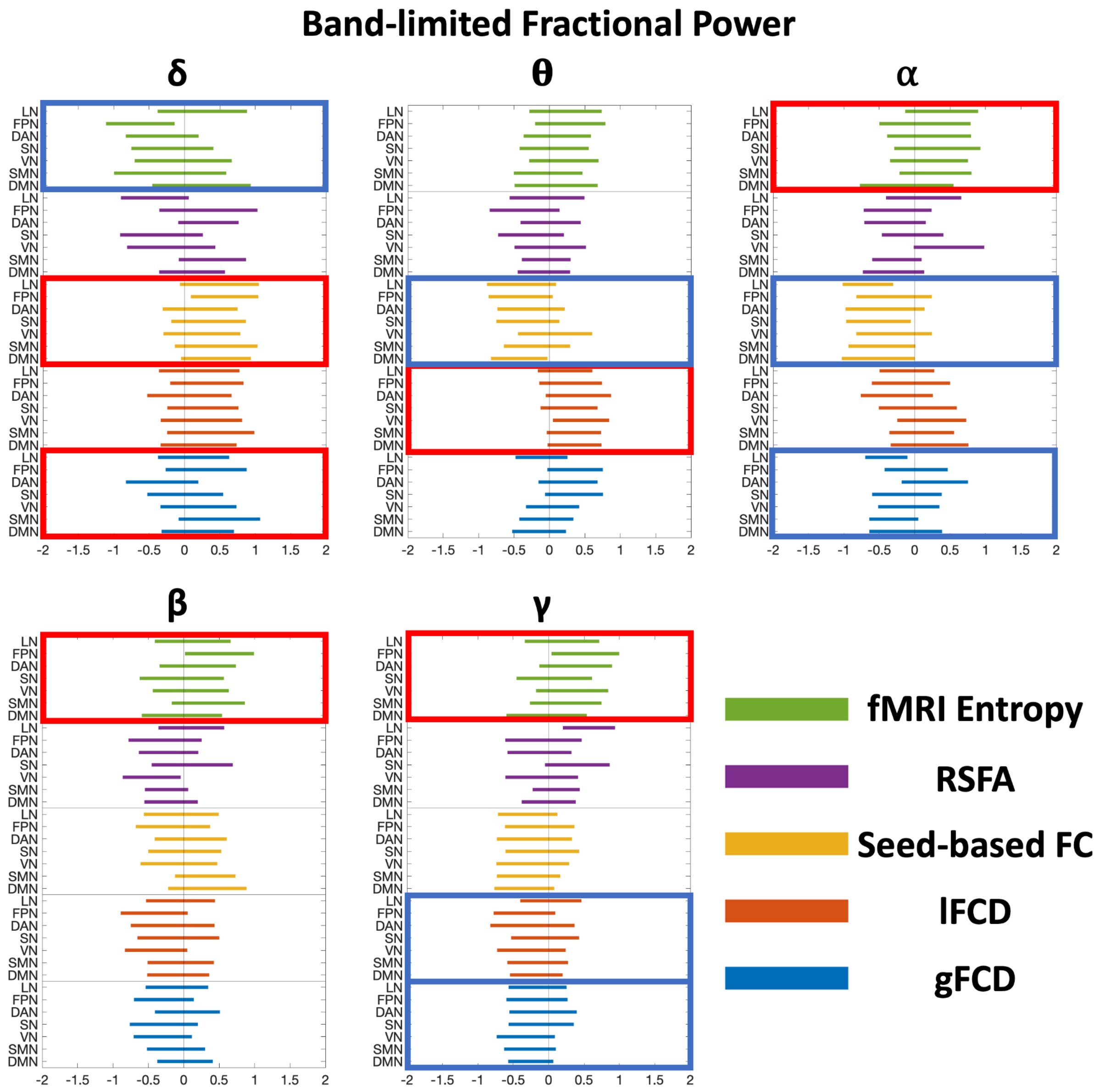


**Figure S6. Effect size of associations between rs-fMRI metrics and EEG band-limited fractional power in each individual network, with macrovascular correction.** Each bar represents a 95% confidence interval of the effect size, and its color corresponds to the rs-fMRI metrics as specified by the legend. The box indicates significant whole-brain associations with the color indicating the direction: red for positive and blue for negative.

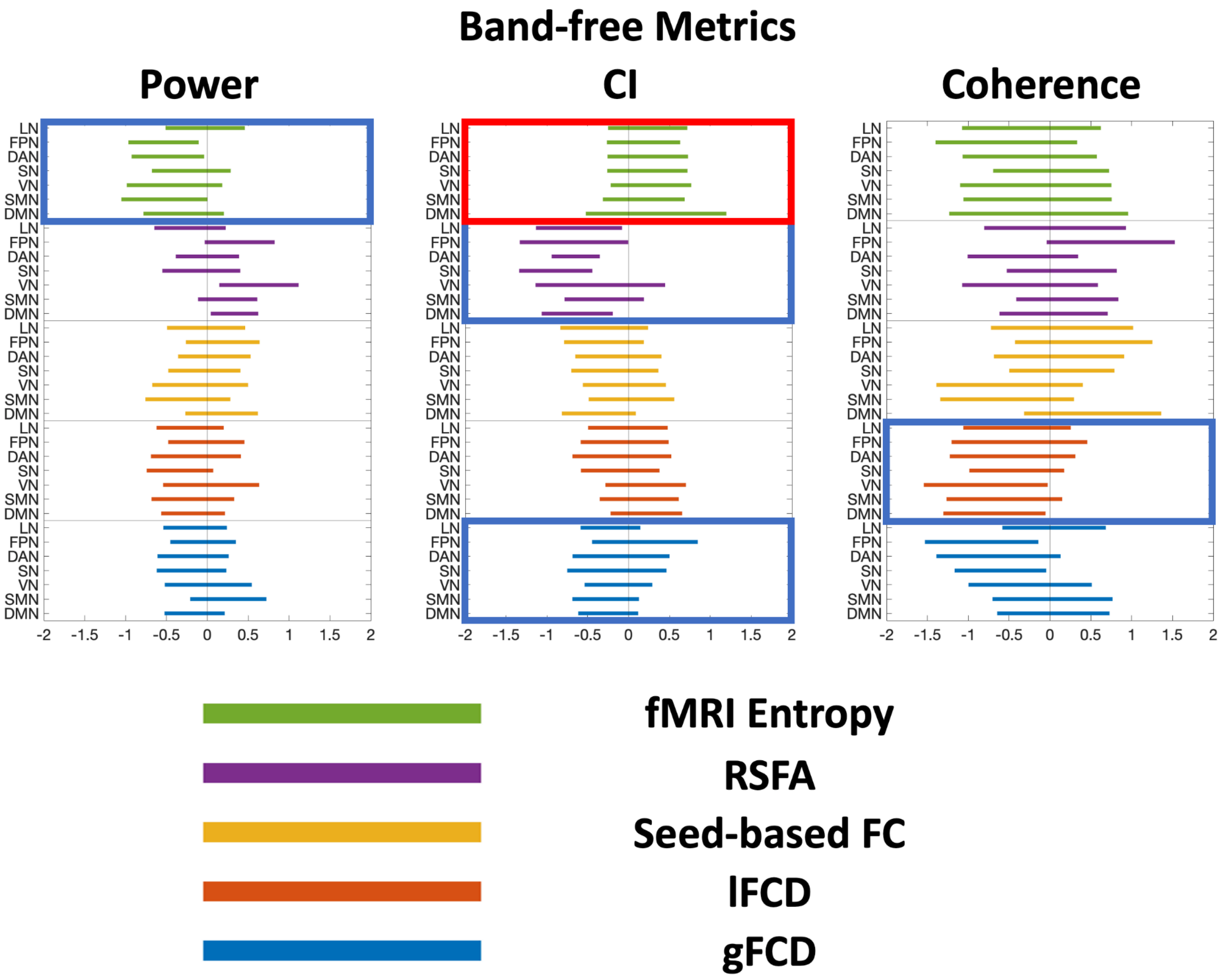


**Figure S7. Effect size of associations between rs-fMRI metrics and EEG band-free metrics in each individual network, with macrovascular correction.** Each bar represents a 95% confidence interval of the effect size, and its color corresponds to the rs-fMRI metrics as specified by the legend. The box indicates significant whole-brain associations with the color indicating the direction: red for positive and blue for negative.
